# A novel biopolymer synergizes type I IFN and IL-1β production through STING

**DOI:** 10.1101/2022.07.22.501157

**Authors:** Ashley R. Hoover, Kaili Liu, Samuel Siu Kit Lam, Chun Fung Wong, Alexandra D. Medcalf, Xiao-Hong Sun, Tomas Hode, Lu Alleruzzo, Coline Furrer, Trisha I. Valerio, Wei R. Chen

## Abstract

N-dihydrogalactochitosan (GC) is developed for inducing immune responses. Synthesized from chitosan and galactose, GC is a new chemical entity that significantly enhances the immune-stimulating properties of its parental material, chitosan, making it a promising therapeutic agent. When used in combination with antigenic material, GC stimulates innate and adaptive antitumor and antiviral immunities. However, the mechanism of GC has not been fully investigated. Herein we demonstrate that GC drives type I IFN production and IFN responses in antigen presenting cells (APCs) and has superior potency compared to its corresponding chitosan. More importantly, GC drives alternative activation of STING leading to inflammatory cell death that enhances dendritic cell (DC) activation, which triggers a variety of nucleic acid sensing pattern recognition receptors (PRRs) and IL-1β production. *In vivo*, GC induced a potent response of type I IFN and upregulated genes associated with STING signaling within the tumor microenvironment (TME). Moreover, intratumoral delivery of GC reduced the numbers of M2-like macrophages residing within the TME, while subsequently increasing the number of DCs. Our findings demonstrate GC’s unique ability to activate STING and stimulate a broad type I IFN response which holds therapeutic promise in generating antitumor and antiviral immunities.

## Introduction

N-dihydrogalactochitosan (GC) is a novel immunoadjuvant that is synthesized using galactose and the partially acetylated glucosamine polymer chitosan. Chitosan, in turn, is derived from chitin, which is largely extracted from the shells of crustaceans, and is a polymer that consists of N-acetylglucosamine(*1*). Chitosan is a nontoxic, biodegradable, biocompatible, natural polysaccharide, currently used in a variety of chemical, biotechnological, pharmacological, and medicinal technologies(*2, 3*). Carroll et al, revealed that using chitosan as a vaccine adjuvant enhanced cellular and humoral mediated immunity and drove a type I interferon (IFN) response through stimulator of interferon genes (STING) signaling(*4*). STING adjuvants, such as cyclic dinucleotides, lead to enhanced humoral and cellular immunity capable of preventing and eliminating intracellular pathogen infections following vaccination and generating protective cellular immunity against tumors, making them highly desirable for stimulating sustained and protective immune responses(*5-8*). However, the cyclic dinucleotides must be administered repeatedly, in high quantities, and/or modified to prevent host cell inactivation(*5, 6, 9*). The fact that chitosan stimulates STING signaling makes it a potential immune stimulant. However, due to its poor solubility at a neutral pH, its poorly understood structure-activity relationship, the potential contributions of endotoxins to its observed activity, and the difficulty to appropriately characterize the polymeric mixture(*10*), the potential of chitosan as a clinically relevant immune stimulant is severely limited.

To address these issues and to significantly increase the type I IFN immune responses, we developed a new chemical entity, N-dihydrogalactochitosan (GC), by attaching galactose to chitosan(*11*). The first clinical GC drug candidate, IP-001(*12*), is synthesized and purified under GMP conditions, which included comprehensive structural characterization and extensive tests for metals, endotoxins and other impurities, further addressing the major challenges in applications of chitosan(*10*). GC has been safely used in early clinical trials for cancer treatment in combination with local tumor ablation(*13, 14*).

We have previously demonstrated the immune stimulating potential of GC in the context of local ablation-based immunotherapy for metastatic cancers, particularly by combining photothermal therapy (PTT) ablation and intratumoral delivery of GC(*13-17*). PTT disrupts tumor homeostasis, induces tumor cell necrosis, and releases tumor-specific antigens (TSAs), while GC overcomes the largely anti-inflammatory tumor microenvironment (TME) and enhances the immune response to the released TSAs. Using RNA sequencing analyses, we examined the immune cells within the TME following intratumoral application of GC and found that GC drove a strong type I interferon (IFN) response in both innate and adaptive tumor-infiltrating leukocytes (TILs). The major immune stimulation in ablation-GC treatment was potentiated by GC(*18, 19*). However, only the combination of ablation and GC was able to achieve sustained tumor regression in both treated tumors and untreated distant metastases(*13-17*).

In addition to the observed antitumor immune responses, our studies on the transcriptomic profiles of immune cells treated with GC revealed a significant upregulation of genes involved in antigen presentation, type I IFN responses, and anti-viral responses(*18, 20*). These results indicate that GC is a potent immunostimulant and can be used as a potential antiviral therapeutic and/or a vaccine adjuvant. The promising therapeutic effect of GC and its potential for future clinical applications warrant an investigation to gain a deeper understanding of its mechanism in triggering immune activation.

In the current work, we show that GC is significantly more potent than its corresponding chitosan starting material in inducing IFNβ and the type I IFN response cytokine Cxcl10 in human THP-1 cells. To further examine the mechanism in which GC induces type I IFN responses, we focused on the activation of dendritic cells (DCs), as GC does not activate non-phagocytic cells. In DCs, GC synergized the productions of type I IFN and IL-1β, both of which are dependent on STING signaling. Interestingly, we discovered STING-initiated lysosomal cell death in response to GC, leading to the activation of caspase-1, and the cleavage of IL-1β and gasdermin D. RNA sequencing analysis revealed that STING played a key role in directing GC-mediated DC activation, producing cytokines, and in stimulating a variety of nucleic acid sensing pattern recognition receptors (PRRs). Furthermore, we found that GC was an active molecule and a STING agonist because it was unable to compromise lysosomal integrity or initiate mitochondrial stress in the absence of STING(*4, 21*). The function of GC in initiating such a broad range of nucleic acid sensing PRRs via inflammatory cell death induction is unique and clinically significant, as most immune stimulants/adjuvants lack this combinatorial capability. Our findings demonstrate that GC can function as an effective immune stimulant for immunotherapy against metastatic cancers and as a potent adjuvant for vaccines against intracellular pathogens.

## Results

### GC enhances activation of THP-1 cells over Chitosan

To compare the similarities and potency of GC with its corresponding chitosan staring material, we stimulated human THP-1 cells, which resemble monocytes and macrophages in morphology and differentiation properties, with chitosan and GC of varying concentrations for 48 hours (Fig. 1, Supplementary Fig. 1). Following the stimulation, we found that GC induced greater amounts of IFNβ production from THP-1 cells compared to chitosan across a range of concentrations (Fig. 1a, 1c, Supplementary Fig. 1a-b). Complementary to type I IFN, CXCL10 production was also significantly higher in the GC-mediated THP-1 cells across a range of concentrations (Fig. 1b, 1d, Supplementary Fig. 1c-d). This suggests that the addition of galactose to chitosan enhances chitosan’s ability to induce type I IFN production. To confirm this finding and to capture the total potency of GC, we graphed the area under the curve (AUC) for the entire range of concentrations (0.5-32μg/ml) for both IFNβ and CXCL10 in the GC and chitosan stimulated THP-1 cells (Fig. 1c-d). AUC analysis of both IFNβ and CXCL10 revealed that GC was significantly more potent than chitosan in stimulating type I IFN and IFN responses in THP-1 cells across a broad range of concentrations (Fig. 1c-d).

**Fig. 1:**
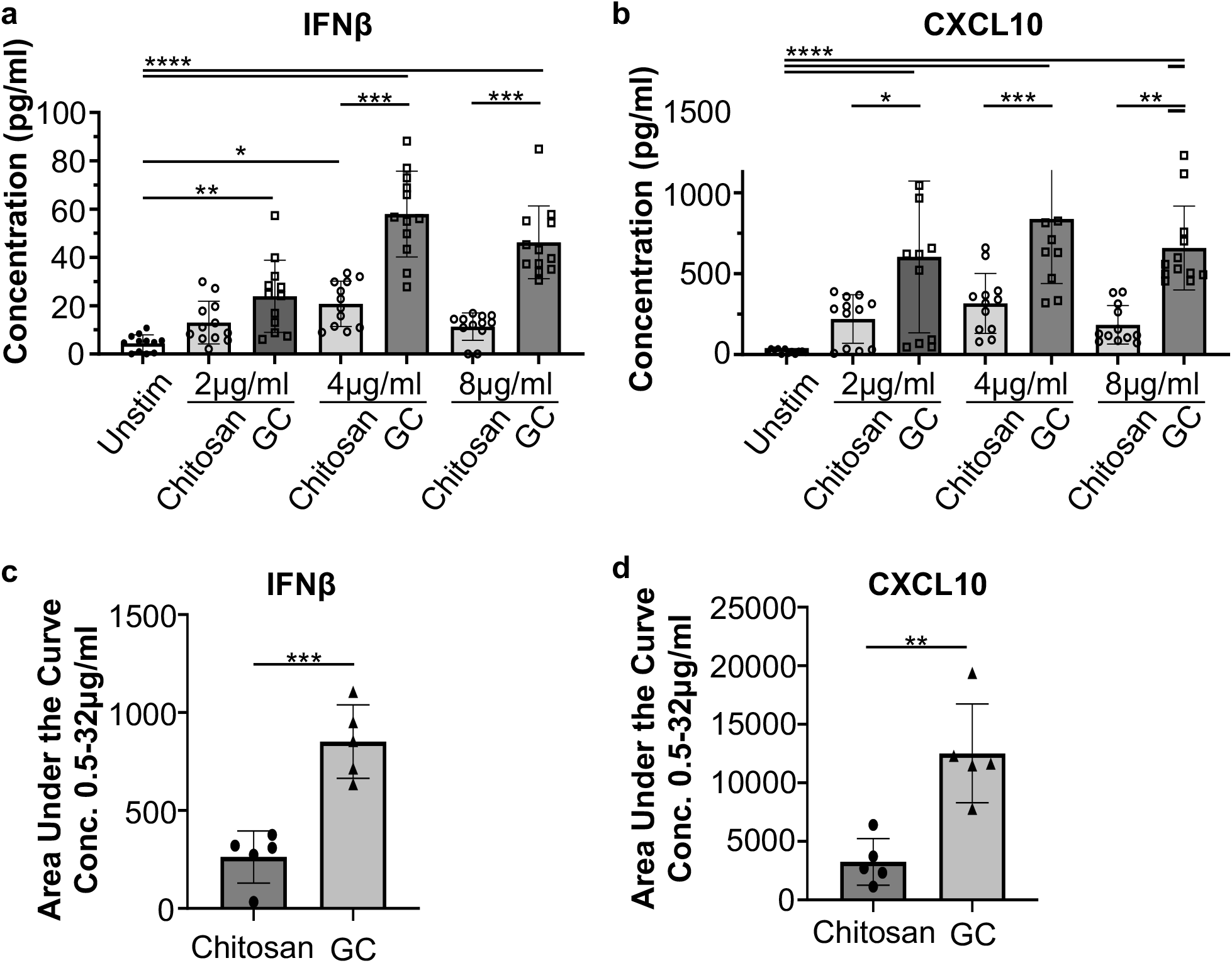
GC induces a stronger type I IFN response in THP-1 cells compared to chitosan. THP-1 cells were stimulated with Chitosan and GC of different concentrations for 48 hours prior to analysis. **a-b**, Production of IFNβ and CXCL10 in the supernatant after stimulation of Chitosan and GC (2μg/ml, 4μg/ml, and 8μg/ml). **c-d**, Total production of IFNβ and CXCL10 in the cell supernatant after stimulation using the area under the curve (AUC) for the entire range of concentrations (0.5-32μg/ml). Data are presented as mean +/-s.e.m. n=2 or more independent experiments. Statistical analysis was performed using one-way analysis of variance (one-way-ANOVA). *p<0.05, **p<0.005, ***p<0.0005, ****p<0.00005.

### GC directly activates unpolarized bone marrow-derived dendritic cells (BMDCs) to produce type I IFNs and IL-1β

Since type I IFN-driven immune responses are critical for antitumor immune responses(*22*), (*23*) and the elimination of viral infections(*24*), and since our *in vitro* experiments have ruled out APC-independent, non-specific T cell activation by GC alone (data not shown), we sought to determine whether GC directly activates DCs to produce type I IFN in a similar manner to the THP-1 cells. GC of varying concentrations was used to stimulate naive bone marrow-derived dendritic cells (BMDCs) to determine the optimal concentration (Fig. 2a). At the concentration of 4μg/ml, the mean fluorescent intensities (MFI) of cell surface molecules CD86, CD40, and MHC-II were increased significantly (Fig. 2b), signaling maturation of the DCs. The production level of IFNβ was comparable between 200ng/ml 2’3-cGAMP (a STING agonist) and 4μg/ml GC (Fig. 2c). In addition, GC induced the production of the proinflammatory cytokine IL-1β (Fig. 2d), which correlates with the upregulation of CD86, CD40, and MHC-II (Fig. 1b). A lower dose of GC (0.8μg/ml) was also able to stimulate the production of IFNβ, but not IL-1β and DC maturation (CD86, CD40, and MHC-II), as shown in Fig. 2a-d. Interestingly, the initiation of BMDC cell death correlated with the production of IL-1β, but not IFNβ, suggesting that the production of the two critical cytokines may be independent of each other (Fig. 2c, d and e).

**Fig. 2:**
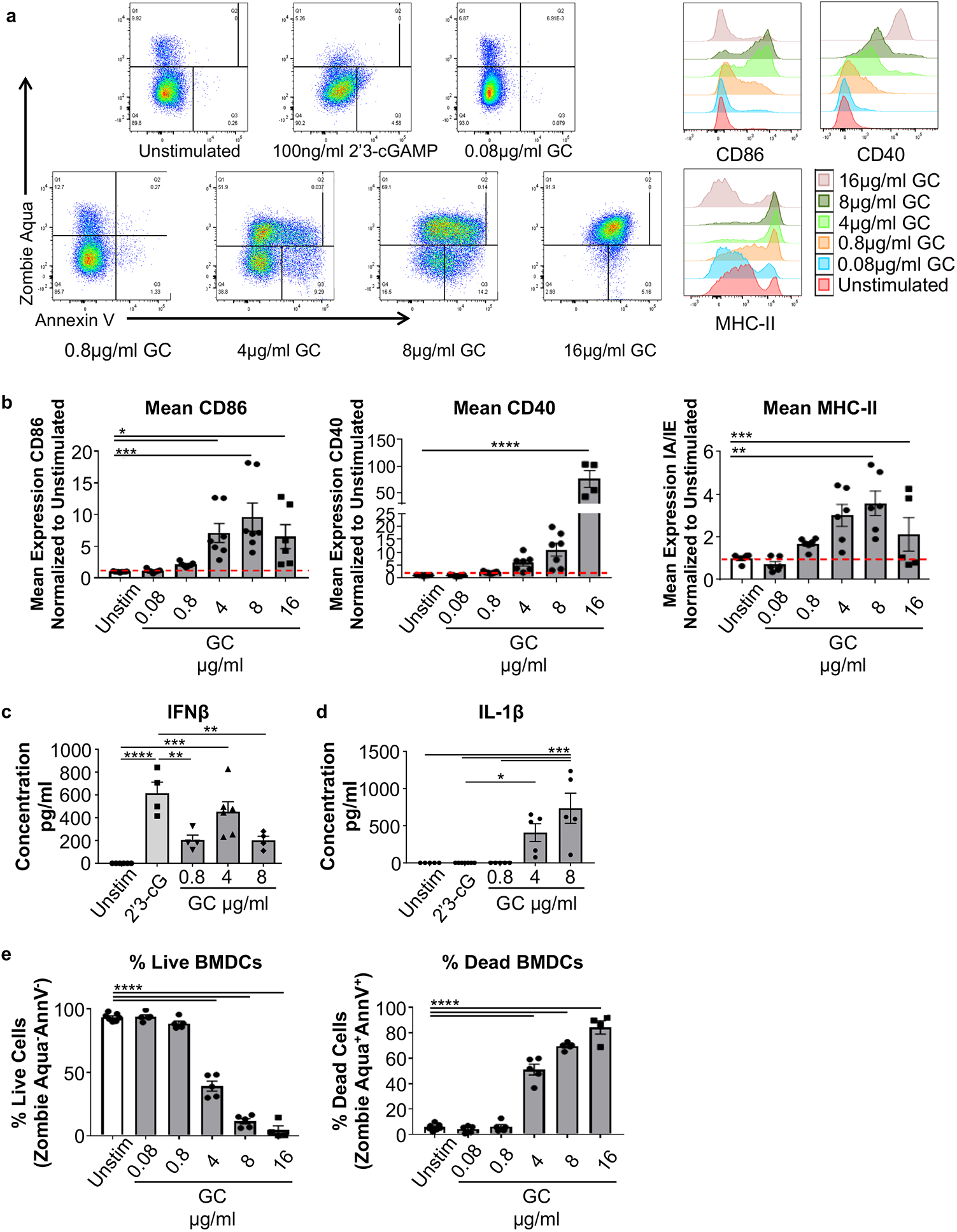
GC directly activates naïve mouse BMDCs and drives the production of Type I IFN and IL-1β. **a**, Mean expressions of CD86, CD40, and MHC-II of live BMDCs after stimulation by GC of various concentrations for 24 hours. Events were pregated on leukocytes/singlets/CD11b^+^CD11c^+^ cells. **b**, Mean expressions of CD86, CD40, and MHC-II, normalized to unstimulated controls and expressed as fold change. Events were pregated on leukocytes/singlets/CD11b^+^CD11c^+/^live cells. **c-d**, Productions of IFNβ and IL-1β cytokines by BMDCs after different stimulations. Supernatant of the BMDC culture was harvested 24 hours after stimulation for ELISA. **e**, Percentages of live (Ghost Dye^-^ Annexin V^-^) and dead (Ghost Dye^+^ Annexin V^+^) cells, pregated as in **a**. Data are presented as mean +/-s.e.m. n=3 or more independent experiments. Statistical analysis was performed using one-way analysis of variance (one-way-ANOVA). *p<0.05, **p<0.005, ***p<0.0005, ****p<0.00005.

To confirm that the productions of these cytokines are independent, we correlated the BMDC cell death and activation of these cytokines. The mean expressions of CD86 and MHC-II increased between 10 and 16 hours while CD40 expression began to increase within 30 minutes of exposure to GC (Supplementary Fig. 2a, b). BMDC cell death began between 10 and 16 hours after GC simulation (Supplementary Fig. 2c). IFNβ production began between 6 and 10 hours, prior to the initiation of BMDC cell death and activation (Supplementary Fig. 2c, d). This correlates with the fact that low doses of GC can induce type I IFN but not cell death, confirming that cell death is not required for type I IFN production (Fig. 2a-d). Interestingly, the production of IL-1β began 16 hours after GC stimulation (Supplementary Fig. 2d), demonstrating that BMDC cell death and the upregulation of CD86 and MHC-II were correlated with the production of IL-1β (Supplementary Fig. 2c, d). This time course revealed interesting kinetics between the production of type I IFN and IL-1β, induced by GC, which warrant further investigation.

### GC induces proinflammatory cell death to drive cytokine production and BMDC activation

To understand the correlation between the GC-mediated BMDC death and activation, as well as IL-1β cytokine production (Supplementary Fig. 1c, d), we explored different inflammatory cell death pathways(*25*). We first investigated pyroptosis, which results from caspase-1/11 cleavage of a pore forming protein gasdermin D(*26*), since the activation of caspase-1 also initiates the cleavage and release of IL-1β(*27*). To determine if GC induced pyroptosis through caspase-1 cleavage, we pre-incubated BMDCs with a caspase-1 inhibitor VX765. VX765 had no effect on IFNβ production, but inhibited IL-1β cleavage (Fig. 3a), as expected. Interestingly, VX765 did not inhibit GC-mediated cell death (Fig. 3b, c), suggesting that the death of BMDCs does not result from caspase-1 activation. Since caspase-11 can also activate gasdermin D and drive pyroptosis(*28, 29*), we observed the gasdermin D levels and found they were significantly reduced by VX765 (Fig. 3d), suggesting that caspase-1, not caspase-11, predominantly drives gasdermin D activation in response to GC. These results reveal that GC-mediated BMDC death was not caused by gasdermin D mediated pyroptosis. Furthermore, BMDC activation is unaltered in the presence of VX765 (Fig. 3e), suggesting that caspase-1 cleavage is not required for GC-induced BMDC activation.

**Fig. 3:**
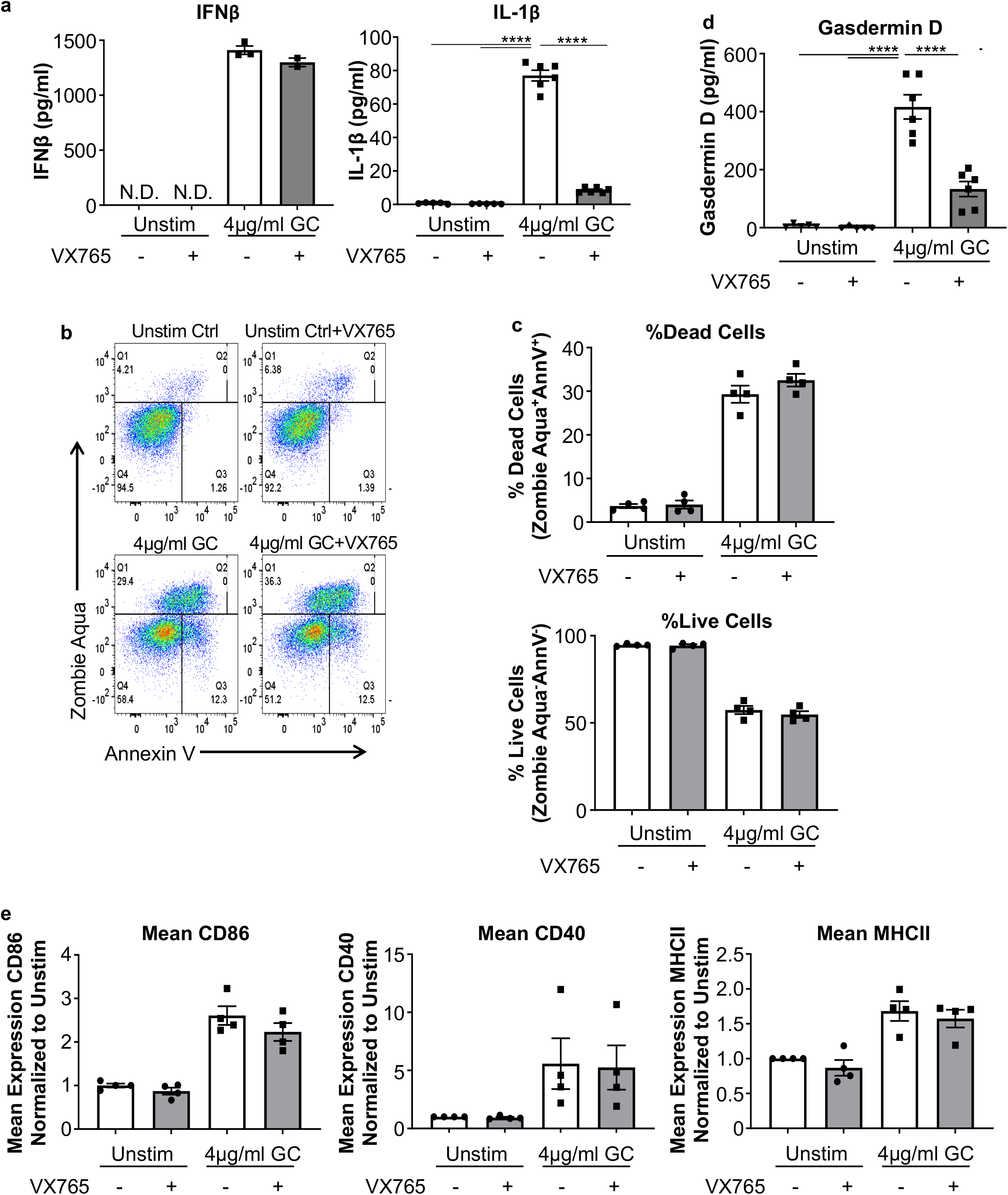
GC drives inflammatory cell death of mouse BMDCs independent of caspase-1 mediated pyroptosis. Mouse BMDCs were pretreated with caspase-1 inhibitor VX-765 for 45 minutes prior to the addition of GC. Cells and supernatants were collected after 24 hours. **a**, Production of IFNβ And IL-1β by BMDCs after GC stimulation for 24 hours. The culture supernatant was used for ELISA. **b**, Visualization of GC mediated cell death using Ghost Dye and Annexin V gating. Events were pregated on leukocytes/singlets/CD11b^+^CD11c^+^ cells. **c**, Percentage of live (Ghost Dye^-^ Annexin V^-^) and dead (Ghost Dye^+^ Annexin V^+^) cells, pregated on leukocytes/singlets/CD11b^+^CD11c^+^ cells. **d**, Expression of Gasdermin D in BMDCs after GC stimulation for 24 hours. Culture supernatant was used for ELISA. **e**, Mean expressions of CD86, CD40, and MHC-II, normalized to unstimulated controls and expressed as fold change. Events were pregated on leukocytes/singlets/CD11b^+^CD11c^+/^Live cells. Data are graphed as mean +/-s.e.m n=3 or more independent experiments. Statistical analysis was performed using one-way analysis of variance (one-way-ANOVA). *p<0.05, **p<0.005, ***p<0.0005, ****p<0.00005.

Since necroptosis is another important form of proinflammatory cell death triggered through the phosphorylation of RIPK3 and MLKL(*30*), we investigated whether necroptosis was involved in GC-mediated cell death. BMDCs were pretreated with a RIPK3 phosphorylation inhibitor GSK’872 prior to GC stimulation. Little to no changes occurred in cell death, activation, or cytokine production in BMDCs (Supplementary Fig. 3), implying that necroptosis is not responsible for GC-mediated BMDC cell death.

To further understand GC-mediated BMDC death, live cell imaging was performed using the mouse DC cell line DC2.4, which behaved similarly to BMDCs (data not shown). After stimulation with FITC-labeled GC for 4-6 hours, we noticed that certain numbers of DCs interacting directly with GC and their neighboring DCs underwent dramatic cellular swelling and eventual rupture, leading either to DC cell death and/or activation of the surrounding DCs (Movie 1). These results suggest GC-induced an inflammatory cell death, distinct from the traditional necroptotic and pyroptotic pathways.

### STING is required for inflammatory lysosome-dependent cell death and cytokine production in GC-stimulated BMDCs

Because the STING pathway has been a critical upstream event to the production of type I IFN for chitosan(*4*), we examined the impact of STING on GC-initiated type I IFN production using the STING “golden ticket” mice(*31*). We stimulated BMDCs from wild type and Tmem173^-/-^ (STING deficient) mice with GC. IFNα/β production was completely abrogated in the Tmem173^-/-^ BMDCs with or without GC stimulation (Fig. 4a), indicating STING is required for type I IFN production. Interestingly, IL-1β production is also blocked in the Tmem173^-/-^ BMDCs with GC stimulation (Fig. 4b). The mean expressions of CD86 and MHC-II were reduced in Tmem173^-/-^ BMDCs compared to wild type (Supplementary Fig. 4a), while the mean CD40 expression was not significantly affected (Supplementary Fig. 4a). Interestingly, cell death in response to GC was also dramatically reduced in Tmem173^-/-^ BMDCs compared to wild type (Fig. 4c), suggesting that STING is also required for GC-mediated cell death.

**Fig. 4:**
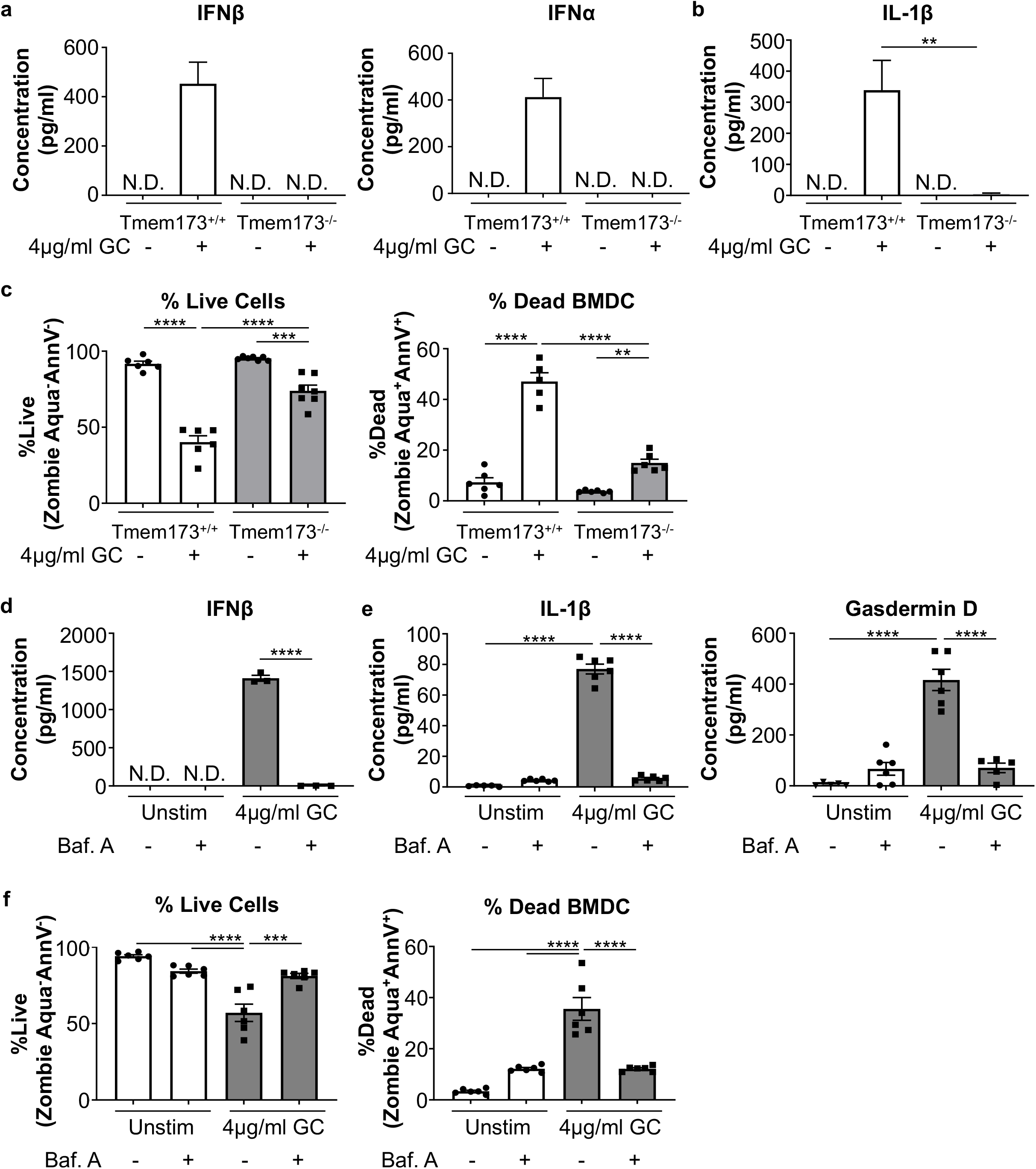
STING and the lysosome are both required for GC to initiate cytokine production and BMDC death. Wild type and Tmem173^-/-^ (STING deficient) BMDCs were stimulated with GC for 24 hours prior to the collection of the supernatant for ELISA and cells for flow cytometry. **a-b**, Production of IFNβ, IFNα, and IL-1β by BMDCs after GC stimulation for 24 hours. The culture supernatant was used for ELISA. **c**, Percentages of live (Ghost Dye^-^ Annexin V^-^) and dead (Ghost Dye^+^ Annexin V^+^) cells, pregated on leukocytes/singlets/CD11b^+^CD11c^+^ cells. **d-e**, Production of IFNβ, IL-1β, and Gasdermin D cytokines by BMDCs, pretreated with lysosomal inhibitor Bafilomycin A for 45 minutes, followed by GC stimulation for 24 hours. Cells and supernatants were collected after 24 hours. Supernatant was for ELISA and cells for flow cytometry. **f**, Percentages of live (Ghost Dye^-^ Annexin V^-^) and dead (Ghost Dye^+^ Annexin V^+^) cells, pregated on leukocytes/singlets/CD11b^+^CD11c^+^ cells. Data are presented as mean +/-s.e.m. n=3 or more independent experiments. Statistical analysis was performed using one-way analysis of variance (one-way-ANOVA). *p<0.05, **p<0.005, ***p<0.0005, ****p<0.00005.

Since recent work by others demonstrated that STING could traffic to the lysosome via an unknown mechanism and initiate lysosomal rupture to stimulate immune cell activation and type I IFN and IL-1β cytokine production(*32*), and since it has been suggested that chitosan has the ability to induce lysosomal permeabilization leading to the production of type I IFN and IL-1β with an unclear mechanism(*21*), we studied the lysosome’s role in GC-mediated cell death and cytokine production. BMDCs were pre-incubated with the lysosomal inhibitor Bafilomycin A (Baf. A), which dramatically inhibited the production of type I IFN, IL-1β, and gasdermin D (Fig. 4d, e). Furthermore, Baf. A significantly reduced GC-mediated BMDC cell death (Fig. 4f), to the level of without GC stimulation. Additionally, the mean expression of cell surface activation markers mimicked that observed on unstimulated Baf. A treated BMDCs (Supplementary Fig. 4b). Collectively, these results indicate that the lysosomes play a key role in GC-mediated cell death, BMDC activation, and cytokine production. Moreover, the results with Baf. A treatment mimicked that observed in the Tmem173^-/-^ BMDCs (Fig. 3a-c versus Fig. 3d-f). This suggests that the interaction between STING and lysosomes is the likely cause of lysosomal mediated cell death, induced by GC, to activate the immune response in a similar manner as described in human monocytes(*33*).

### GC induces robust type I IFN antitumoral and antiviral responses in BMDCs through cellular necrosis

To further investigate the roles of STING in mediating GC-induced cellular responses, we followed the orthogonal experimental design of a two-conditions (without and with GC treatment) and two-genotypes (wide type and Tmem173^-/-^), with interaction terms(*34*) (Supplementary Fig. 5a), to perform bulk RNA sequencing (RNAseq) for BMDCs.

Using the mRNA of protein coding genes, we generated heatmaps of sample-to-sample distances using the variance stabilizing transformation (vsd) values (Fig. 5a). Figure 5a showed that the largest distance existed between the WT BMDCs with and without GC stimulation, suggesting that GC induced broad transcriptional profile changes. Meanwhile, distance between unstimulated WT and Tmem173^-/-^ BMDCs is the smallest, indicating that they both had similar gene expressions without GC stimulation. The responses of Tmem173^-/-^ BMDCs, when stimulated with GC, were remarkably similar to that of the unstimulated wild type and Tmem173^-/-^ BMDCs, but significantly different from that of GC-mediated WT BMDCs, suggesting that STING may play a crucial role in mediating GC activation of BMDCs. We also conducted the differentially expressed genes (DEGs) analysis by comparing transcriptional profiles in GC-treated WT BMDCs versus untreated BMDCs. Expressions of these GC-triggered DEGs shown by the heatmap indicated that GC-promoting genes (upregulated DEGs) and GC-inhibiting genes (downregulated DEGs) contain both STING-dependent and STING-independent transcriptional signatures (Supplemental Fig. 5b).

**Fig. 5:**
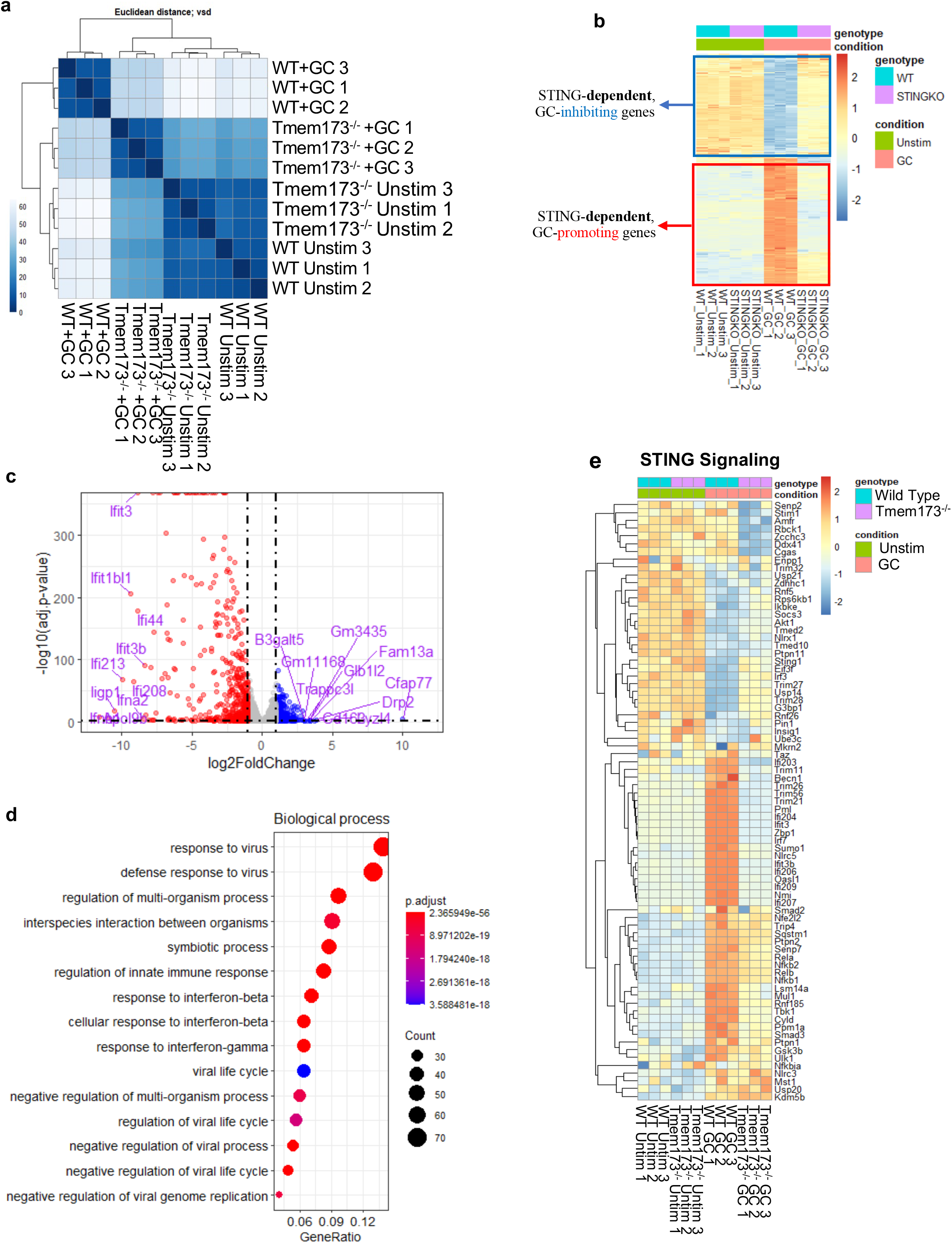
GC is unable to activate BMDCs in the absence of STING. Wild type and STING deficient (Tmem173^-/-^) BMDCs were stimulated with GC for 24 hours prior to collection. Cells were sorted on Live/CD11b^+^CD11c^+^ BMDCs prior to total RNA isolation. **a**, Heatmap of sample-to-sample distances using the variance stabilizing transformation (vsd) values of mRNA expression of protein coding genes. **b**, Heatmap schematically showing the STING-dependent GC-promoting genes (in red box) and GC-inhibiting genes (in blue box). Each row represents a gene (gene name not shown). **c**, Volcano plot demonstrating the expressions of STING-dependent, GC-promoting genes (red color) and GC-inhibiting genes (blue color). Each dot represents a STING-dependent gene. **d**, Dot plot demonstrating the enrichment analysis for STING-dependent GC-promoting genes using biological process (BP) of gene ontology (GO) database. **e**, Heatmap showing the expressions of STING pathway genes in WT and Tmem173^-/-^ BMDCs with and without GC treatment. Wild type and STING deficient (Tmem173^-/-^) BMDCs were stimulated with GC for 24 hours prior to collection.

Next, we dissected the STING-dependent, GC-promoting and -inhibiting transcriptional profiles by using *DESeq2* package(*35*) following the two-factor analysis and visualized in heatmap and volcano plot (Fig. 5b-c). The STING-dependent, GC-induced DEGs contained both downregulated and upregulated expression patterns when compared to WT BMDCs with GC treatment, corresponding to GC-promoting genes (Fig. 5c, left) and GC-inhibitory genes (Fig. 5c, right). As expected, the top GC-promoting genes WT I IFNs (*Ifnb1*) and type I IFN response genes (Fig. 5c, left), consistent with the observation using cytokine ELISA (Fig. 4a).

Our Gene Ontology (GO)(*36*) enrichment analysis also showed that STING-dependent, GC-promoting genes were enriched in biological processes including responses to virus and IFNβ/γ (Fig. 5d). This was further confirmed by our Kyoto Encyclopedia of Genes and Genomes (KEGG)(*37*) enrichment analysis, which showed the involvement of pathways including infection, virus, Toll-like receptor, antigen processing and presentation, necroptosis, RIG-I-like receptor, and TNF signaling (Supplemental Fig. 5c). On the other hand, the STING-dependent, GC-inhibitory genes were enriched in the biological processes and pathways of cell cycle and cell nuclear division (Supplemental Fig. 5d). For example, expressions of cell cycle related genes, including *Ccnb1, Top2a, E2f7, Cdc25c*, and *Mki67*, were significantly downregulated in GC-stimulated WT BMDCs. However, this STING-dependent GC-inhibition was attenuated in Tmem173^-/-^ BMDCs (Fig. 5e). These results support our previous observation that GC treatment led to STING-dependent cell survival/cycle reduction and cell death augmentation (Fig. 4c).

### GC requires STING to activate BMDCs and to promote the activation of a variety of nucleic acid pattern recognition receptors

To further explore the antitumor and antiviral response pathways activated by GC, we focused on positive and negative regulators of the nucleic acid sensing pattern recognition receptors (PRRs) such as STING, MAVS/RIG-I, TL3, and TLR7/9 (Fig. 5e, Supplementary Fig. 6a-c). As expected, the unstimulated samples, regardless of genotype, were nearly identical in their gene expression patterns (Fig. 5e). Furthermore, the genes expressed in the GC-stimulated Tmem173^-/-^ BMDCs were like the unstimulated controls, confirming that STING pathway genes were not activated by GC in its knockout genetic background (Fig. 5e).

In addition, STING signaling can activate TBK1/IRF3 or TBK1/IRF7 signaling pathways and/or canonical and noncanonical NFkB signaling(*38-41*). Based on gene expression patterns in the WT BMDCs, 24 hours after stimulation, GC drove STING signaling preferentially through TBK1/IRF7, but not TBK1/IRF3, in addition to both canonical and noncanonical NFkB signaling (Fig. 5e). Next, we examined the expression of upstream mediators of STING activation: cGAS, Zbp1 (also known as DAI), DDX41, and IFI16 (also known as Ifi204). The expressions of cGAS and DDX41 were equivalent between the unstimulated and GC-mediated WT BMDCs but were decreased in GC-stimulated Tmem173^-/-^ BMDCs (Fig. 5e). Surprisingly, Zbp1 and Ifi204 are specifically enriched in the GC-stimulated WT BMDCs. Both are cytosolic nucleic acid sensing proteins that signal through STING to initiate type I IFN production. Zbp1 is unique in that it is capable of also binding dsDNA and dsRNA(*42*), ribonucleoprotein complexes(*43*), and also activating RIPK3, to drive necroptosis(*44*). This is complemented by our *in vitro* data demonstrating that blocking RIPK3 activation slightly reduced GC-mediated BMDC cell death (Supplementary Fig. 3b). Since Ifi204 is a cytosolic dsDNA sensor that plays a key role in herpes simplex virus type 1 (HSV-1) immunity(*45*) and it was preferentially induced in response to GC as opposed to cGAS (Fig. 5e), our results support our hypothesis that, in contrast to chitosan(*4*), GC activation of STING in DCs may be cGAS independent.

Since it has been suggested that chitosan is a mitochondrial toxin and that the production of ROS and the release of mitochondrial DNA/RNA activate an immune response(*4*), and in that case, other nucleic acid PRRs and the NLRP3 inflammasome should still be activated even in the absence of STING, we generated heatmaps of genes involved in RIG-I/MDA-5, TLR-3, -7, and -9 signaling. The results revealed that these pathways were only activated in the GC-mediated WT BMDCs, not in the Tmem173^-/-^ or unstimulated BMDCs (Supplementary Fig. 6a-c). This suggests that mitochondrial function was not directly affected by GC. Moreover, these data showed that the activation of RIG-I/MDA-5, and TLR-3, -7, and -9 signaling pathways were downstream of STING. The activation of these pathways is consistent with the notion that GC is mediating a form of pro-inflammatory cell necrosis, specifically through STING, to activate neighboring BMDCs through a variety of nucleic acid sensing pathways. This unique method of DC activation, combined with the fact that GC upregulates an antiviral response, suggests that GC could function as an effective antiviral vaccine adjuvant.

When WT BMDCs was compared to the Tmem173^-/-^ BMDCs upon GC stimulation, no stress response genes were upregulated in the GC-mediated Tmem173^-/-^ BMDCs, indicating that GC was not a direct mitochondrial toxin and/or it did not result in direct lysosomal leakage, as genes for lysophagy and mitophagy were not upregulated in the absence of STING compared to WT BMDCs (Supplementary Fig. 7a, b).

### Intratumoral injection of GC drives a type I IFN, proinflammatory tumor microenvironment

To determine how GC exerts its effects on myeloid cells *in vivo*, we injected MMTV-PyMT tumors with GC, isolated the tumor infiltrating immune cells, and performed single-cell RNA-Sequencing (scRNAseq) as previously described(*18*). To explore the heterogeneity of the myeloid cells, we used shared-nearest neighbor (SNN) to cluster the myeloid cells. Using principal component analysis (PCA), we obtained 12 clusters (C0-C12), annotated using a series of commonly used immune cell genes (Fig. 6a and Supplementary Fig. 8a). The proportions of immune cells in the 12 clusters were compared between the untreated and GC-mediated cells (Fig. 6b). GC enhanced the proportions of G-MDSC (+281%), monocytes (C9, +50%), macrophages (C1, +107%), plasmacytoid DCs (pDCs, C6, +53.1%), conventional DCs (C11 and C12, +77.6% and +54.3, respectively), and granulocytic monocyte derived suppressor cells (G-MDSCs, C2, +281%). Conversely, GC decreased the proportion of select macrophages (C0, - 53.6%; C4, -143%; C5, -12.9%; C7, -73.4%; C8, -87.4%; and C10, -71.4%), within the TME (Fig. 6b, Supplementary Fig. 8b).

Since STING can interact with *Tbk1, Ikbkg*, and *Traf6* to drive type I IFN production, and canonical or noncanonical NFkB signaling(*38, 40, 45*), we examined several genes involved in STING signaling cascades and discovered the most significant upregulation of genes occurred in the monocytes and the three DC clusters (Figure 6c). Monocytes with upregulated expressions of *Tbk1, Ikbke (IKK*ε*), Irf3 Ikbkg (NEMO), Ikbkb (IKK*β*), Chuk (IKK*α), and *Nfkb1 (p50*), correspond to the conventional STING signaling, type I IFN production, and the activation of canonical NFkB signaling gene targets(*38*). A similar pattern was observed in cDCs (C12), and pDCs (C6). In C11, cDCs with upregulated *Tbk1, Nfkb1, Irf7*, and *Irf3*, suggest conventional STING signaling. Interestingly, cells in C11 also have upregulated genes *Nfkb2* and *Relb*, both involved in noncanonical NFkB signaling(*40*). These results indicate that STING signaling is in fact occurring in the monocytes and DC subtypes in response to GC, corroborating our *in vitro* findings using BMDCs. Interestingly, the macrophages did not respond to GC in a similar manner as DCs, a phenomenon we have also observed *in vitro* (data not shown).

**Fig. 6:**
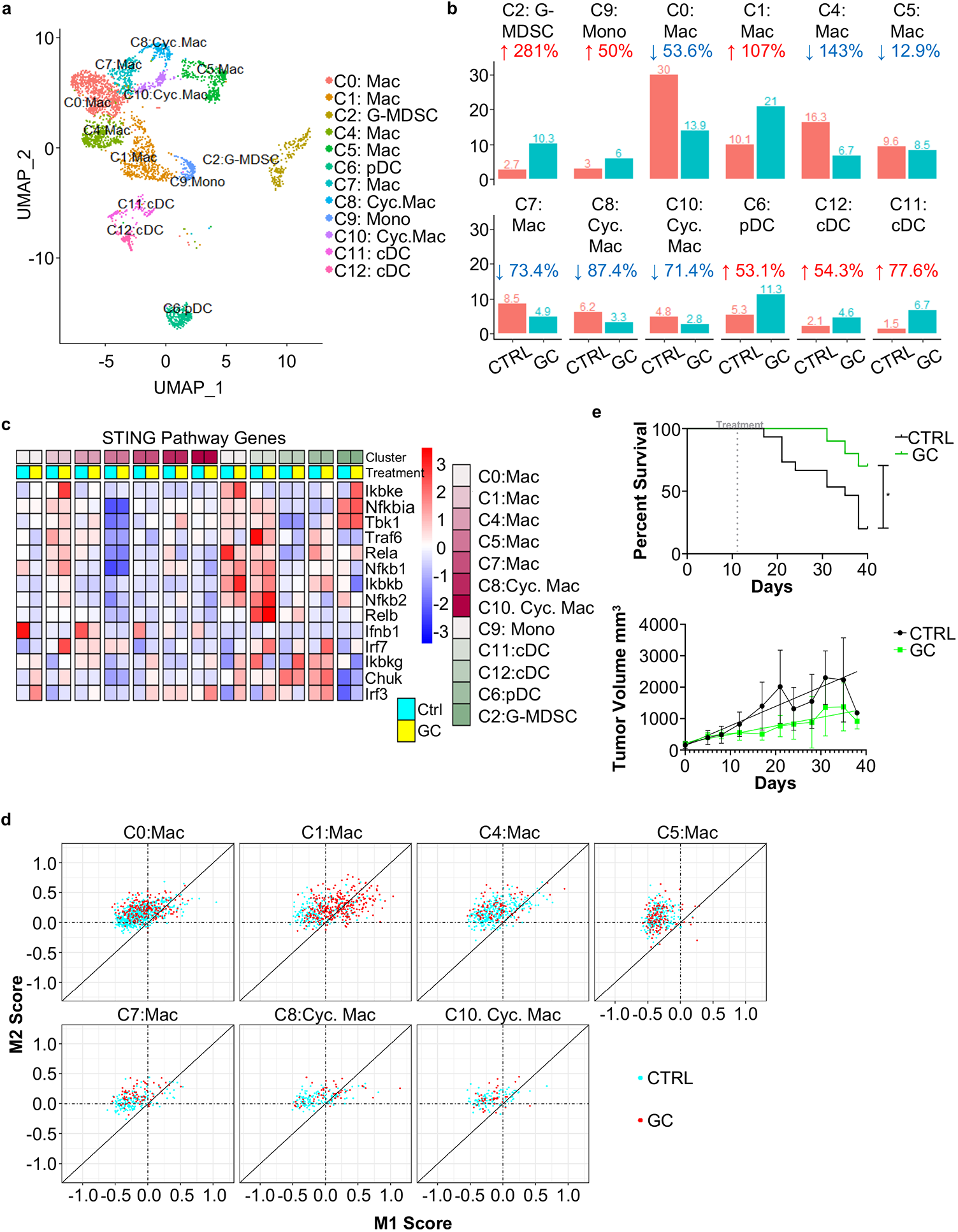
GC remodels myeloid cell proportions/heterogeneity and activation status in the tumor microenvironment. Wild type FVB females were injected with MMTV-PyMT tumor cells directly into the mammary tissue. Once the tumors reached ∼0.5 cm^3^, animals were either left untreated or injected with 1% GC (0.1 ml). Tumors were collected 10 days post treatment, and tumor infiltrating CD45^+^ cells were purified and used for single cell RNA-sequencing. For survival studies, the tumor-bearing animals were monitored for 30 days. **a**, UMAP of myeloid cell subtypes, including monocyte, macrophage (Mac), dendritic cell (DC), myeloid-derived suppressing cell (MDSC). **b**, Bar plots displaying the proportions of myeloid cells from untreated (CTRL) and GC treated tumors. Fold change from CTRL to GC was displayed. **c**, Heatmap highlighting expression of genes involved in the STING signaling pathway. **d**, Scatter plots highlighting the macrophage clusters and their M1- and M2-like macrophage score. **e**, Survival of mice bearing MMTV-PyMT tumor and tumor growth following intratumoral GC injection. Statistical significance was performed using log-rank test. *: p < 0.05

We next examined the expression of cytokines, chemokines, and regulatory molecules in all the myeloid cell subsets. Interestingly, the monocytes and pDCs did not express proinflammatory cytokines while cDC expressed upregulated proinflammatory cytokines and chemokines (Supplementary Fig. 8c). This is significant for T cell activation in the TME as DCs play a major role in activating and cross priming T cells in the TME(*46*). Examining the macrophage clusters more closely revealed that the only macrophage cluster that increased in numbers, C1 (Fig. 6b), has the upregulated expressions of *Cd40, Il1b, Cxcl10, Tgm2*, and *Nos2*, and molecules and cytokines associated with inflammation (Supplementary Fig. 8c). The other macrophage subsets, which decreased in number with the addition of GC, exhibited increases in anti-inflammatory and immune regulating molecules, such as *Il10, Tgfb, Arg1, Vegfa, Apoe*, and *Socs3*, to various degrees (Supplementary Fig. 8c). These results suggest that the macrophages in C1 exhibit a more inflammatory phenotype, while the remaining macrophage clusters are more anti-inflammatory.

To further explore the inflammatory (M1-like) and anti-inflammatory (M2-like) macrophage phenotypes present in the TME, we used a scatter plot to compare traditional M1- and M2-like gene expression patterns for untreated control cells (shown in red in Fig. 6d) and GC-treated cells (shown in turquoise in Fig. 6d). Macrophages in clusters 0, 4, 5, 7, 8, and 10 revealed a similar M2-like protumor signature between the untreated and GC-treated cells, albeit there were fewer numbers in the GC-treated group (Fig. 6d). Interestingly, the macrophages in C1 demonstrated that a portion of cells with upregulated genes associated with M1-like macrophages, suggesting that these cells are more inflammatory (Fig. 6d). More importantly, the macrophages that shifted towards the M1-like phenotype were almost exclusively from the GC-treated tumors. This corresponds well with the activation signature observed in the GC-treated macrophages in C1 (Supplementary Fig. 8c). While GC does not appear to repolarize M2-like macrophages towards an M1-like phenotype, it does significantly reduce their numbers, likely through inflammatory cell death similar to GC-mediated DC death.

Since it is known that tumor associated macrophages (TAMs) are indicators of treatment resistance, tumor progression, and metastases(*47-49*), we assessed survival of mice bearing a syngeneic MMTV-PyMT tumor after intratumoral injection of GC (1%, 0.1ml), once the tumors reach 0.5mm^3^. Surprisingly, a single GC injection significantly slowed tumor growth and prolonged survival (Fig. 6e). These results suggest that GC is a potent immune stimulant that can activate DCs and macrophages in the TME to potentiate antitumor immunities.

## Discussion

In the current work we investigated the mechanism by which GC activates strong antitumor and antiviral immune responses. First, we discovered that GC could not directly activate non-phagocytic cells such as T cells or tumors cells (data no shown), consistent with the previous results using chitosan for activation of APCs(*21*). Instead, GC must be actively engulfed to activate APCs such as DCs. Work by others also suggested that, following phagocytosis, chitosan triggers lysosomal leakage, leading to the activation of the NLRP3 inflammasome/pyroptosis pathway and mitochondrial stress, resulting in the activation of the cGAS/STING pathway to produce type I IFNs(*4, 21*). These studies concluded that chitosan was passive in inducing the immune response and that immune stimulation occurred via lysosomal leakage-induced cellular stress.

We synthesized N-dihydrogalactochitosan (GC) to create a new molecule that is both more soluble and significantly more potent than its corresponding chitosan, allowing it to become a viable drug candidate for clinical use. Using human THP-1 cells, which resemble monocytes and macrophages in morphology and differentiation properties, we showed that GC was much more effective than chitosan in inducing type I IFN production and activation (Fig. 1). We also found that GC induced cell death that correlates with the production of IL-1β (Fig. 2). We determined that GC did not induce pyroptosis (Fig. 3c); instead, it initiated STING-dependent lysosomal cell death (Fig. 4c). Without STING, BMDC death was drastically reduced and type I IFNs and IL-1β were completely abrogated (Fig. 4a, b). Furthermore, we ascertained that endosome/lysosome fusion played a key role in GC-initiated production of type I IFN and IL-1β (Fig. 4d).

In human monocytes, it has been reported that in certain viral infections, cGAS activation of STING can cause STING trafficking to the lysosome via an unknown mechanism and it initiates lysosomal rupture to activate the NLRP3 inflammasome(*33*). The mechanism of GC appears to be similar, but the involvement of cGAS was not clear. We compared the activation of WT and Tmem173^-/-^ BMDCs following GC stimulation and found that Tmem173^-/-^ BMDCs stimulated with GC were similar to the unstimulated controls (Fig. 4). The results in Fig. 4 indicate that STING is required for DC activation and GC was not a mitochondrial toxin; it did not cause significant amounts of lysosomal leakage in the absence of STING. Furthermore, Tmem173^-/-^ BMDCs stimulated with GC did not elevate genes involved in mitochondrial or lysosomal stress (Supplemental Fig. 7), suggesting that GC plays an active role in stimulating STING and that STING is directly required for GC-mediated activation and lysosomal leakage. This is corroborated by the fact that cGAS expression is not upregulated in response to GC (Fig. 5e). It should be noted that other STING activating molecules, such as Zbp1 and IFI16 (Ifi204), were highly enriched by GC (Fig. 5e). Whether these molecules are required for GC-mediated STING activation requires further investigation.

It has recently been demonstrated that STING can be activated by a polyvalent polymer containing a seven membered ring with a tertiary amine (PC7A)(*50*). This molecule does not bind STING at the same site as the cyclic dinucleotides like 2’3-cGAMP, and this alternate binding to STING actually prolongs the type I IFN response compared to cyclic dinucleotides(*50*). Furthermore, it was found that cGAMP initiated a strong immune response within the first couple of hours, peaking at 6 hours, while PC7A initiated a peak stimulation between 24 and 48 hours, effectively sustaining the STING activation(*50*). Our kinetic studies with GC revealed that type I IFN production did not begin until ∼10 hours after stimulation and continued to increase over a 24-hour period (Supplementary Fig. 2d). This suggests that GC may act similarly to PC7A in sustaining STING signaling. However, the advantage of GC, comparing to PC7A, is its capability in driving alternative activation of STING, resulting in an inflammatory cell death that further enhances APC recruitment and activation through a variety of pattern recognition receptors (PRRs), such as toll-like receptors (TLR), nucleotide-binding and oligomerization domain (NOD)-like receptors (NLRs), and retinoic acid-inducible gene I (RIG-I)-like receptors (RLRs), as shown in Supplemental Fig. 6.

Our analysis of the myeloid cells isolated from GC-injected MMTV-PyMT tumors revealed a STING activation signature in cDCs and a dramatic shift in the macrophage populations (Fig. 6b-c). Furthermore, GC increased the proportions of DCs, while simultaneously decreasing the anti-inflammatory macrophages residing within the TME (Fig. 6b, d). Moreover, the simple addition of GC into the TME significantly reduced tumor growth and prolonged survival (Fig. 6e). These results indicate that GC is a strong type I IFN stimulant that can alter the TME, making it more inflammatory which precedes immune mediated tumor killing.

In summary, GC triggered conventional and alternative activations of STING to synergize type I IFN and IL-1β productions. These novel functions and the unique mechanism make GC a promising therapeutic immunostimulant/immunoadjuvant in generating antitumor and antiviral immunities.

## Materials and methods

### GC and chitosan

N-dihydrogalactochitosan (GC) and chitosan was provided by Immunophotonics, Inc. (St. Louis, MO). GC was synthesized, purified, characterized, and tested for impurities under GMP conditions, and the same batch of GC was used in all experiments. The chitosan used for experiments shown in Figure 1 and Supplemental Figure 1 was from the same batch of chitosan that was used as starting material in the synthesis of the GC. Additionally, the chitosan was processed and purified under identical conditions as the GC (except for the synthetic steps) and was characterized and tested to ensure that the chitosan and GC materials were identical, except for the galactose moieties on the GC molecule.

### Bone Marrow-Derived Dendritic Cell Cultures

Bone marrow was isolated from 8 to 12-week-old male or female mice. Briefly, the animals were euthanized according IACUC approved protocols. The hind legs were removed and separated from the muscles and sinew. The heads of the bones were removed and 1x PBS was pushed through the bones using a 25-gauge needle and a 1ml syringe. The red blood cells were lysed, and cell were resuspended in RPMI containing 20% heat inactivated fetal calf serum, pen/strep, and β-mercaptoethanol. About 5×10^6^ bone marrow cells were plated into 100mm non-tissue culture treated plates containing 20ng/ml GMCSF (Biolegend, Cat# 640920). On day 3 5ml of fresh media containing 20ng/ml of GMCSF was added to the cultures. On day 6 non-adherent cells were collected and medium was removed and replaced with fresh RPMI containing 20ng/ml GMCSF and placed back into the 100mm plate. On day 7 the nonadherent BMDCs were collected for stimulation.

### BMDC Activation

BMDCs were stimulated with GC or 2’3-cGAMP (InvivoGen, Cat# tlrl-nacga23-1) for 0-24 hours prior to ELISA or flow cytometry. About 10^6^ cells/ml BMDCs were plated into a non-tissue culture treated 96 well round bottom plate and then stimulated with 0.08-16μg/ml GC. For 2’3-cGAMP stimulation, 200ng of 2’3-cGAMP was first encapsulated into viromer green transfection reagent (Origene, Cat# TT100301) according to the manufacturer’s instructions prior to incubation with BMDCs.

VX765 was purchased from InvivoGen (Cat# inh-vx765i-1) and used at a concentration of 10μM. BMDCs were pre-incubated for 30 minutes with VX765 prior to the addition of GC. Bafilomycin A was purchased from InvivoGen (Cat# tlrl-baf1) and used at a concentration of 100nM. BMDCs were pre-incubated for 30 minutes with bafilomycin A prior to the addition of GC. GSK’872 was purchased from Millipore Sigma (Cat# 530389) and used at a concentration of 3μM. BMDCs were pre-incubated for 45 minutes with GSK’872 prior to the addition of GC.

### RNA-Sequencing

BMDCs were cultured for 24 hours, then harvested and stained with ghost dye BV510 for viability and CD11b APC-Cy7 and CD11c FITC. Live CD11b+CD11c+ BMDCs were sorted on the BD FACS ARIA. RNA was isolated from the sorted BMDCs using the Quick-RNA microprep kit purchased from Zymo Research (Cat# R1050). RNA was isolated according to the manufacturer’s protocol. For sequencing the 20M reads mRNA prep and sequencing service was performed for preparation of the RNA for NovaSeq PE150 reads on the NovaSeq6000.

### Bioinformatics analysis (Quality control, read trimming, mapping to genome, and identification of DEGs)

A quality check for the raw sequencing data was conducted using *FastQC* (v0.11.9) to detect common issues in RNA-Seq data. The reads were then trimmed with *Trimmomatic* (v0.39) to remove low quality bases(*51*). The quality of the reads was re-evaluated with *FastQC* after this step to validate the quality improvements. The RNA-seq reads from each sample were mapped to the mouse mm10 genome assembly using the *HISAT2. Samtools* was used to manipulate the *HISAT2* generated SAM files into BAM files(*52*). *FeatureCounts* program from *Subread* package was used to count mapped RNAseq reads for genomic features(*53*). *DESeq2* was used for differential expressed genes (DEG) analysis based on the negative binomial distribution. The resulting P-values were adjusted using the Benjamini and Hochberg’s approach for controlling the false discovery rate. Genes with an adjusted P-value (P adj) < 0.05 as determined by *DESeq2* were assigned as differentially expressed. Gene ontology (GO) analysis of DEG was performed using *clusterProfiler*(*54*).

### Flow Cytometry of BMDCs

All samples were run on the BD FACSCelesta and data were analyzed using FlowJo v10.6.1. After BMDC stimulation for 0-24 hours with GC or 2’3-cGAMP, BMDCs were isolated and stained with the antibodies, CD11c-FITC (Tonbo Bioscience, cat# 35-0114), CD11b-APC-Cy7 (Tonbo Bioscience, Cat# 25-0112), CD40-PE-Cy7 (Biolegend, Cat# 124622), MHCII-RedFluor710 (Tonbo Biosciences, Cat# 80-5321), CD86-BV605 (Biolegend, Cat# 105037), Ghost Dye-Violet-510 (Tonbo Biosciences, Cat# 13-0870), and Annexin V-APC (Biolegend, Cat# 640920). Briefly, cells were stained on ice with all antibodies and viability dye for 20 minutes. The cells were then washed and resuspend in Annexin V staining buffer for 15 minutes and then immediately ran on the BD FACSCelesta. Briefly, cells were gated on CD11c^+^CD11b^+^ cells and then assessed for ghost dye BV510 and Annexin V expression. For the mean expression of CD40, CD86, and MHCII, the cells were gated on CD11c^+^CD11b^+^ live cells before analysis. Bar graphs were generated using Graphpad prism software.

### Cytokine ELISA

After BMDC stimulation with GC or 2’3-cGAMP for 0-24 hours, BMDCs were isolated and pelleted, and the supernatant was collected and frozen at -80ºC until the ELISAs were performed. The mouse IFNβ and IFNα ELISA kit was purchased from R&D systems (Cat# MIFNB0), mouse IL1β ELISA kits were purchased from ThermoFisher Scientific (Cat# 88-7013-22), and mouse gasdermin D ELISA kits were purchased from IBL America (Cat# IB99570). ELISAs were performed according to manufacturers’ instructions. The ELISA plates were read using the Biotek Synergy H1 Hybrid plate reader, and the data were analyzed and graphed using the Graphpad Prism software.

### Animals

All animal studies were either approved by the IACUCs of Oklahoma Medical Research Foundation and the University of Oklahoma. C57BL/6, C57BL/6J-*Sting1*^*gt*^*/*J, and FVB mice were purchased from Jackson Laboratories (Stock number: 000664, 017537, and 001800).

### Syngeneic tumor cell transplantation

MMTV-PyMT murine breast tumor organoids were isolated from FVB/N-Tg (MMTV-PyVT) 634Mul/J mice as previously described(*55*). Briefly, cells were incubated overnight in mammary epithelial cell media (DMEM/F12 supplemented with 10% fetal bovine serum (Sigma Aldrich, F2442-500ML) 100 U/mL penicillin-streptomycin (Sigma Aldrich, P0781-20×100ml), 5ug/mL insulin-transferrin-selenium (ThermoFisher Scientific, 51500056), 1ug/mL hydrocortisone (Sigma Aldrich, H0888-10G), 10ng/mL mouse epidermal growth factor (EGF) (ThermoFisher Scientific, 53003018), and 50 ug/mL gentamicin (Genesee Scientific, 25-533). Cells were washed twice with Hepes Buffered Saline (HBSS) (Gibco, 14175-095) trypsinized, and resuspended to a concentration of 1×10^5^ cell per 20uL. Cells were injected into the mammary fat pad of wild type FVB mice without clearing. The incision was closed with Vetbond tissue adhesive (3M). Once the tumors reached 0.5cm^3^ the tumors were either left untreated or injected with 100μl of 1% GC (Immunophotonics, St. Louis, MO).

### Sample preparation and single-cell RNA sequencing library generation

In each of the two treatment groups, immune cells from four mice were used for single-cell RNA sequencing. Nine days after treatment, tumor tissues were isolated, minced with scalpels, and digested with Collagenase IV and Dnase I at 37°C for 20-30 minutes. After enzymatic digestion, immune cells were enriched using lymphocyte separation medium. The enriched cells were then subjected to magnetic bead separation (EasySep Mouse Streptavidin Rapidspheres Isolation Kit, Stem Cell, 19860) to remove the EpCAM^+^ cells (CD326 1:200, ThermoFisher Scientific, 13-5791-82). The EpCAM-depleted cells were stained with antibodies against CD45 and a viability dye. Live CD45^+^ cells (CD45 1:100, Biolegend, 103112), were sorted using MoFlo and then processed for droplet-based 3’ end scRNAseq by encapsulating sorted live CD45^+^ tumor-infiltrating immune cells into droplets via a 10× Genomics platform according to the manufacturer’s instructions (10× Genomics). Paired-end RNA-seq was performed via an Illumina NovaSeq 6000 sequencing system.

### ScRNA-seq Library Generation

Droplet-based 3’ end scRNA-seq was performed by encapsulating sorted live CD45^+^ tumor-infiltrating immune cells into droplets using the 10x Genomics platform. cDNA libraries were prepared using Chromium Single Cell 3’ Reagent Kits v3 according to the manufacturer’s protocol (10x Genomics). The generated scRNA-seq libraries were sequenced using an Illumina NovaSeq 6000.

### Alignment, Barcode Assignment, UMI Counting

The Cell Ranger pipeline was used to conduct sample demultiplexing, barcode processing, and single-cell 3’ counting. The Linux command *cellranger mkfastq* was applied to demultiplex raw base call (BCL) files from the illumine NovaSeq6000 sequencer, into sample-specific fastq files. Then, fastq files for each sample were processed with *cellranger count*, which was used to align samples to mm10 genome, filter and quantify.

### Data Preprocessing with Seurat R Package

Seurat-guided analyses (https://satijalab.org/seurat/vignettes.html) were used to preprocess and integrate datasets from different treatment groups(*56*). Genes that were expressed in less than 5 cells or cells that expressed less than 8000 and more than 6000 genes were excluded. Also, cells that expressed less than 512 and more than 92600 counts or with a mitochondria percentage over 10% were excluded. Most variable genes were identified using the FindVariableFeatures function by setting feature numbers as 2000. Principal component analysis (PCA) was performed on the scaled matrix (with most variable genes only) using the first 30 principal components (PCs). Both tSNE and UMAP dimensional reductions were carried out using the first 20 PCs to obtain two-dimensional representations of the cell states. For clustering, we used the function FindClusters that implements a shared nearest neighbor (SNN) modularity optimization-based clustering algorithm on 30 PCs with resolution 0.5 for default analysis.

### Identification of Cluster-specific Genes and Marker-based Classification

Cells clusters were obtained by using the FindClusters function of the Seurat R package, which identifies clusters through an SNN modularity optimization–based algorithm. The biological cell type identities of each cluster were annotated with the assistance of an immune-cell scoring algorithm comparing the differentially expressed gene (DEG) signatures obtained from Seurat with the Immunological Genome Project Database (ImmGen)(*57*). This *in silico* cell type annotation was verified by using known canonical immune cell genes.

### Gene Set Enrichment Analysis

Differential Gene Expression was obtained by using the FindMarkers function in Seurat. MAST was used as the methodology. Ranked gene lists were then analyzed for gene set enrichment by using the clusterProfiler R package (*58*). Hallmark gene sets (H) from the Molecular Signature Database (MSigDB, https://www.gsea-msigdb.org/gsea/msigdb) were used in these analyses (*59*).

### Statistical Analysis

Evaluations for tumor growth and FACS data were conducted using one-way ANOVA. MAST test was used for analyzing differential gene expression in selected cell clusters. P values less than or equal to 0.05 were considered statistically significant throughout (*, p ≤ 0.05; **, p ≤ 0.01).

## Data and Software Availability

The accession number for the scRNAseq data reported in this paper is GEO: GSE150675 (https://www.ncbi.nlm.nih.gov/geo/query/acc.cgi?acc=GSE150675). Analysis of such data can also be available upon request.

## Acknowledgement

This work was supported in part by the National Cancer Institute (R01CA205348 and R01CA269897), and the Oklahoma Center for the Advancement of Science and Technology (HR16-085 and HF20-019).

## Ethics declarations

WRC is co-founder and a member of the Board of Directors of Immunophotonics, Inc. CFW, TH, SSKL, and LA declare a conflict of interest as employees with minority ownership stakes of Immunophotonics, Inc., the manufacturer of the proprietary immunostimulant GC.

