## Supplementary Figure Legends for "A novel biopolymer synergizes type I IFN and IL-1β production through STING"

**Supplementary Fig. 1: GC induces a stronger type I IFN response in THP-1 cells compared to chitosan.**

THP-1 cells were stimulated with Chitosan and GC of different concentrations for 48 hours prior to analysis. **a-b,** Production of IFNβ in the supernatant after stimulation of Chitosan and GC (0.5μg/ml, 16μg/ml, and 32μg/ml). **c-d,** Production of d CXCL10 in the supernatant after stimulation of Chitosan and GC (0.5μg/ml, 16μg/ml, and 32μg/ml). Data are presented as mean +/- s.e.m. n=2 or more independent experiments. Statistical analysis was performed using one-way analysis of variance (one-way-ANOVA). *p<0.05, **p<0.005, ***p<0.0005, ****p<0.00005.

**Supplementary Fig. 2: GC activation of BMDCs is a prolonged process over a period of 24 hours.**

Cell viability, activation, and cytokine production of wild type BMDCs stimulated with GC (4μg/ml) over a course of 24 hours. Supernatants of BMDC after stimulation were harvested at different time points. **a**, Cell viability over a course of 24 hours visualized by the expression of Ghost Dye and Annexin V. Events were pregated on leukocytes/singlets/CD11b^+^CD11c^+^ cells. **b**, Mean expressions of CD86, CD40, and MHC-II, normalized to unstimulated controls and expressed as fold change. Events were pregated on leukocytes/singlets/CD11b^+^CD11c^+/^Live cells. **c**, The percentages of live cells (Ghost Dye^-^ Annexin V^-^ and Ghost Dye^-^ Annexin V^+^), and the percentage of dead cells (Ghost Dye^+^ Annexin V^+^), pregated on leukocytes/singlets/CD11b^+^CD11c^+^ cells. **d**, Production of IFNβ and IL-1β cytokines from GC-stimulated BMDCs. Supernatants of GC stimulated BMDCs were collected at various time points for ELISA. Data are presented as mean +/- s.e.m. n=3 or more independent experiments. Statistical analysis was performed using one-way analysis of variance (one-way-ANOVA). *p<0.05, **p<0.005, ***p<0.0005, ****p<0.00005.

**Supplementary Fig. 3: GC-mediated cell death is independent of necroptosis.**

Mouse BMDCs were pretreated with RIPK3 phosphorylation inhibitor GSK’872 for 45 minutes prior to the addition of GC. Cells and supernatants were collected after 24 hours, with 2’3-cGAMP as a positive control. **a**, Mean expressions of CD86, CD40, and MHC-II, normalized to unstimulated controls and expressed as fold change. Events were pregated on leukocytes/singlets/CD11b^+^CD11c^+^/Live cells. **b**, Percentages of live (Ghost Dye^-^ Annexin V^-^) and dead (Ghost Dye^+^ Annexin V^+^) cells, pregated on leukocytes/singlets/CD11b^+^CD11c^+^ cells. **c**, Production of IFNβ and IL-1β cytokines from GC-stimulated BMDCs. Supernatants of BMDCs after stimulation were collected and used for ELISA. Data are presented as mean +/- s.e.m. n=3 or more independent experiments. Statistical analysis was performed using one-way analysis of variance (one-way-ANOVA). *p<0.05, **p<0.005, ***p<0.0005, ****p<0.00005.

**Supplementary Fig. 4: Blocking STING or lysosomal activity prevents the activation of BMDCs by GC.**

**a,** Mean expressions of CD86, CD40, and MHC-II from wild type and STING deficient (Tmem173^-/-^) BMDCs, normalized to unstimulated controls and expressed as fold change. Wild type and STING deficient BMDCs were stimulated with GC and cells were collected 24 hours later. Events were pregated on leukocytes/singlets/CD11b^+^CD11c^+^/Live cells. **b,** Mean expressions of CD86, CD40, and MHC-II from wild type and lysosomal inhibited (Bafilomycin A) BMDCs, normalized to unstimulated controls and expressed as fold change. Wild type and lysosomal inhibited BMDCs were stimulated with GC and cells were collected 24 hours later. Events were pregated on leukocytes/singlets/CD11b^+^CD11c^+^/Live cells. Data are presented as mean +/- s.e.m. n=3 or more independent experiments. Statistical analysis was performed using one-way analysis of variance (one-way-ANOVA). *p<0.05, **p<0.005, ***p<0.0005, ****p<0.00005.

**Supplementary Fig. 5: STING deficient BMDCs stimulated with GC are enriched in housekeeping genes.**

Wild type and STING deficient (Tmem173^-/-^) BMDCs were stimulated with GC for 24 hours prior to collection. Cells were sorted on Live/CD11b^+^CD11c^+^ BMDCs prior to total RNA isolation. **a**, Schematic of a two-factor (genotype, treatment) orthogonal experiment design with 4 groups of 12 samples. **b**, STING-dependent/-independent, GC-promoting/-inhibiting genes (gene names not shown). GC-promoting genes were upregulated differentially expressed genes (DEGs) from comparison of GC stimulation versus no stimulation in WT BMDCs. GC-inhibiting genes were downregulated DEGs from comparison of GC stimulation versus no stimulation in WT BMDCs. **c**, Dot plot demonstrating the enrichment analysis for STING-dependent, GC-promoting genes using Kyoto Encyclopedia of Genes and Genomes (KEGG) database. **d**, Dot plot demonstrating the enrichment analyses for STING-dependent, GC-inhibiting genes using GO and KEGG databases.

**Supplementary Fig. 6: GC activates a variety of nucleic acid sensing signaling pathways, dependent of STING signaling.**

Wild type and STING deficient (Tmem173^-/-^) BMDCs were stimulated with GC for 24 hours prior to collection. Cells were sorted on Live/CD11b^+^CD11c^+^ BMDCs prior to total RNA isolation. Heatmaps were generated to visualize the expression of genes involved in activation of MAVS/RIG-I (**a**), TLR3 (**b**), and TLR7/9 (**c**) signaling pathways, comparing unstimulated wild type, STING-deficient BMDCs with GC-stimulated wild type, STING-deficient BMDCs.

**Supplementary Fig. 7: Upregulation of lysophagy and autophagy associated genes in GC stimulated BMDCs require STING signaling**

Wild type and STING deficient (Tmem173^-/-^) BMDCs were stimulated with GC for 24 hours prior to collection. Cells were sorted on Live/CD11b^+^CD11c^+^ BMDCs prior to total RNA isolation. Heatmaps were generated to visualize the expression of genes involved in the activation of lysophagy (**a**) and mitophagy (**b**), comparing GC-stimulated wild type and STING-deficient BMDCs.

**Supplementary Fig. 8: Tumor infiltrating myeloid cells respond positively to intratumoral GC injection.**

Wild type FVB female mice were injected with MMTV-PyMT tumor cells directly into the mammary tissue. Once the tumors reached ~0.5cm^3^, animals were either left untreated or injected with 1% GC (0.1ml). Tumors were collected 10 days post treatment, and tumor infiltrating CD45^+^ were purified and used for single cell RNA-sequencing. **a**, Myeloid cell populations were labeled based on the expression of several genes enriched in each subset. **b**, UMAP of resolution 0.5 demonstrating the commonality and differences between the infiltrating immune cells in untreated and GC-treated tumors. **c**, Heatmap to visualize the expression of several genes involved in myeloid cell activation and inflammatory functions.
