## Supplementary Figures for "A novel biopolymer synergizes type I IFN and IL-1β production through STING"

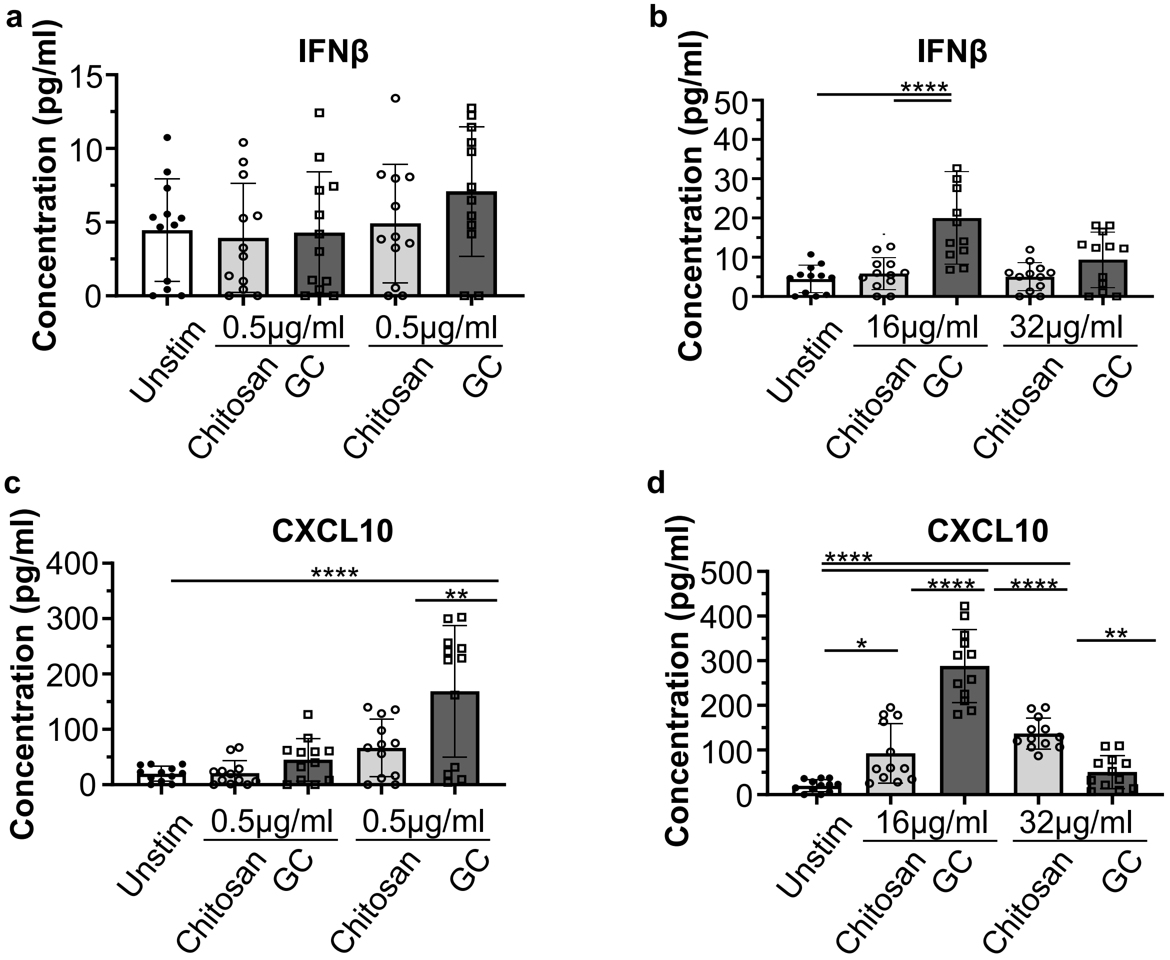
a
IFNβ
Concentration (pg/ml)
15
10
5
0
Unstim
0.5µg/ml
Chitosan
GC
0.5µg/ml
Chitosan
GC
b
IFNβ
Concentration (pg/ml)
50
40
30
20
10
0
****
Unstim
16µg/ml
Chitosan
GC
32µg/ml
Chitosan
GC
c
CXCL10
Concentration (pg/ml)
400
300
200
100
0
****
**
Unstim
0.5µg/ml
Chitosan
GC
0.5µg/ml
Chitosan
GC
d
CXCL10
Concentration (pg/ml)
500
400
300
200
100
0
****
****
****
*
**
Unstim
16µg/ml
Chitosan
GC
32µg/ml
Chitosan
GC

**Figure S1**

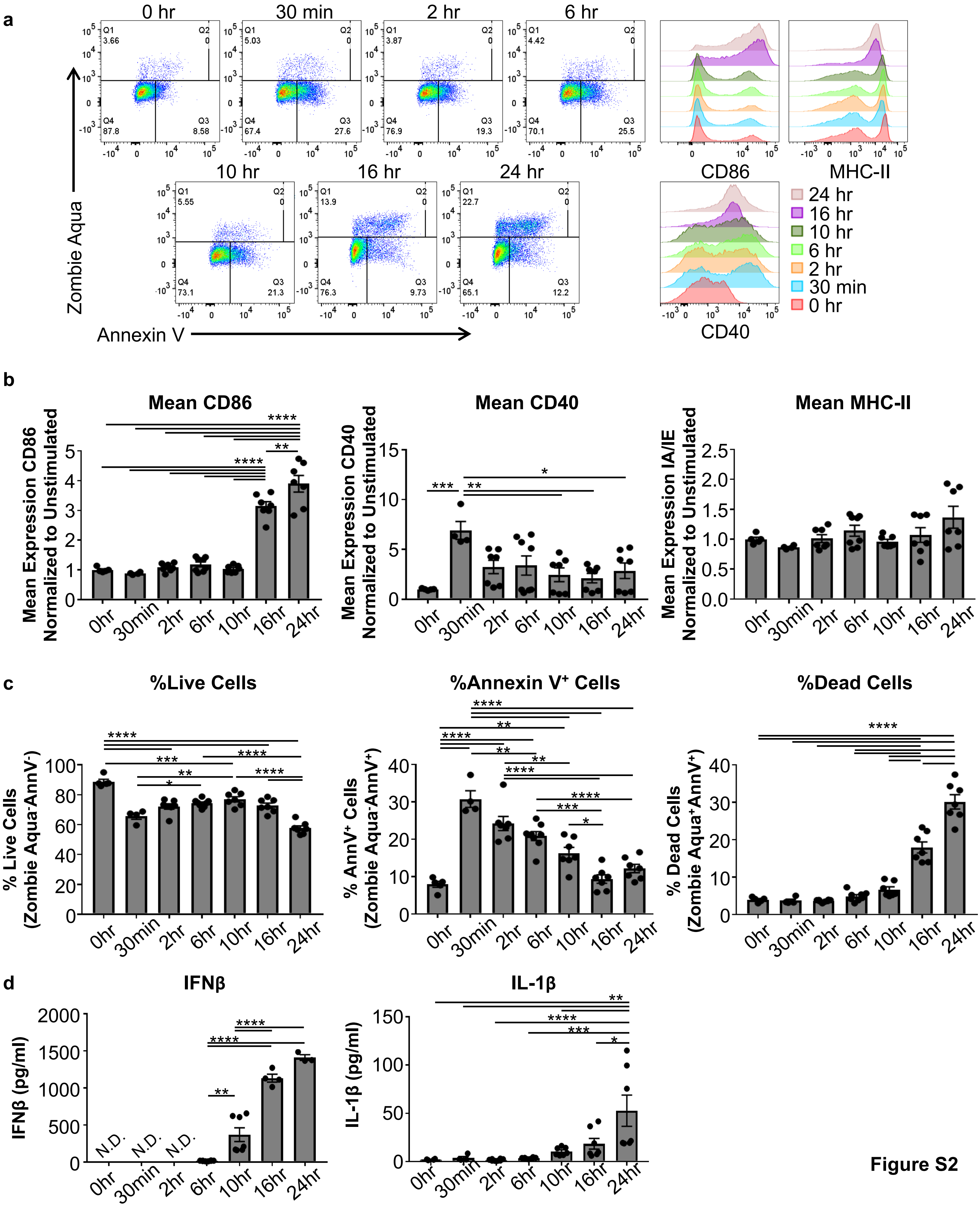
a
0 hr
30 min
2 hr
6 hr
10 hr
16 hr
24 hr
Zombie Aqua
Annexin V
CD86
MHC-II
CD40
24 hr
16 hr
10 hr
6 hr
2 hr
30 min
0 hr
b
Mean CD86
Mean CD40
Mean MHC-II
Mean Expression CD86 Normalized to Unstimulated
Mean Expression CD40 Normalized to Unstimulated
Mean Expression IA/IE Normalized to Unstimulated
c
%Live Cells
%Annexin V+ Cells
%Dead Cells
% Live Cells (Zombie Aqua-AnnV-)
% AnnV+ Cells (Zombie Aqua-AnnV+)
% Dead Cells (Zombie Aqua+AnnV+)
d
IFNβ
IL-1β
IFNβ (pg/ml)
IL-1β (pg/ml)
N.D.
0hr
30min
2hr
6hr
10hr
16hr
24hr

Figure S2

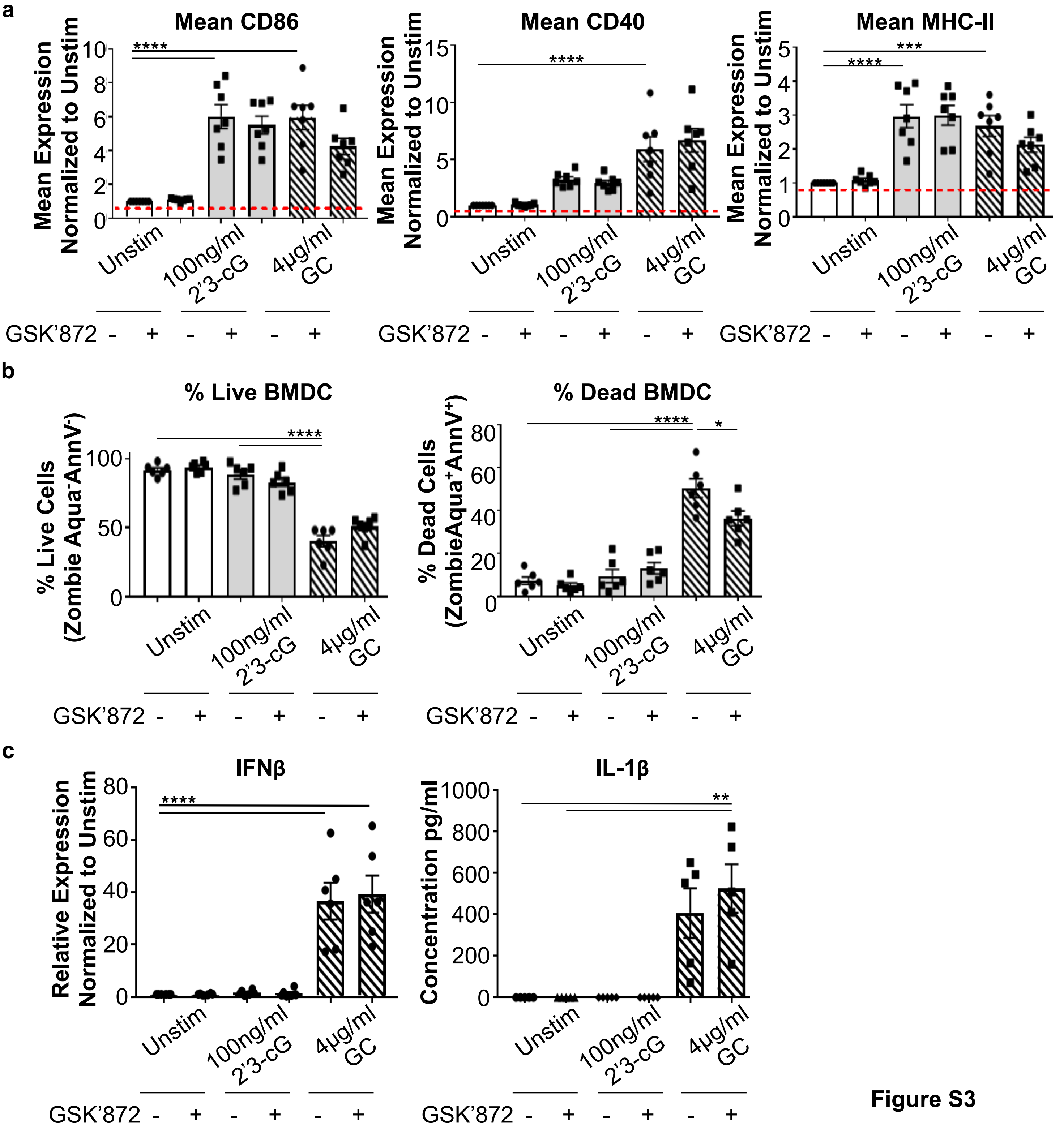
a
Mean CD86
Mean Expression Normalized to Unstim
****
Mean CD40
****
Mean MHC-II
****
***
b
% Live BMDC
% Live Cells (Zombie Aqua⁻AnnV⁻)
****
% Dead BMDC
% Dead Cells (ZombieAqua⁺AnnV⁺)
****
*
c
IFNβ
Relative Expression Normalized to Unstim
****
IL-1β
Concentration pg/ml
**
Unstim
100ng/ml 2'3-cG
4μg/ml GC
GSK'872 - + - + - +

Figure S3

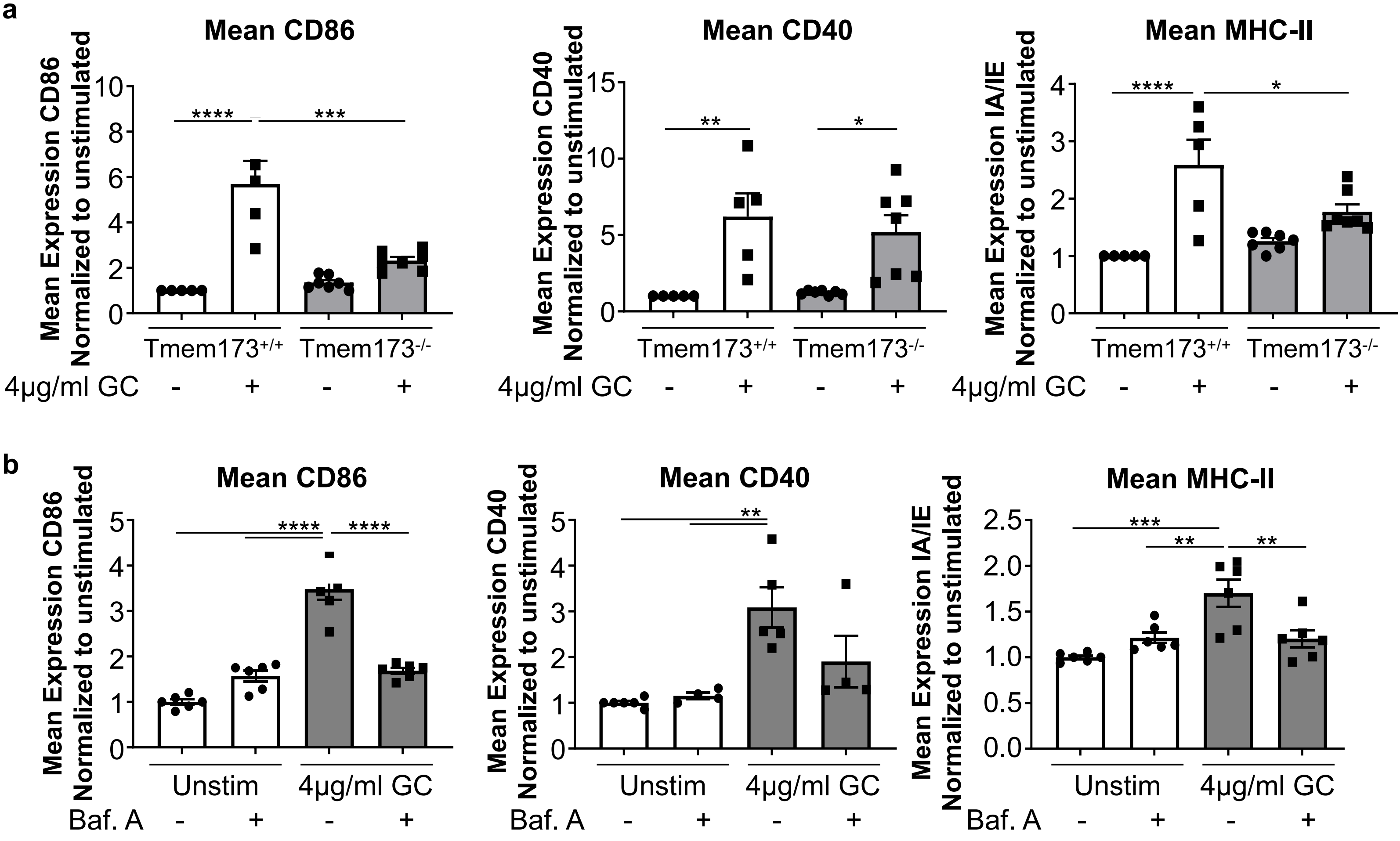
a
Mean CD86
Mean Expression CD86 Normalized to unstimulated
****
***
Tmem173+/+
Tmem173-/-
4μg/ml GC
Mean CD40
Mean Expression CD40 Normalized to unstimulated
**
*
Mean MHC-II
Mean Expression IA/IE Normalized to unstimulated
****
*
b
Mean CD86
****
****
Unstim
4μg/ml GC
Baf. A
Mean CD40
**
Mean MHC-II
***
**
**

Figure S4

**a**

| | | **treatment** | |
|---|---|---|---|
| | | **without GC** | **with GC** |
| **genotype** | **WT** | Group **A**: 1, 2, 3 | Group **C**:7, 8, 9 |
| | **Tmem173$^{-/-}$ (STING KO)** | Group **B**: 4, 5, 6 | Group **D**: 10,11, 12 |

**b**

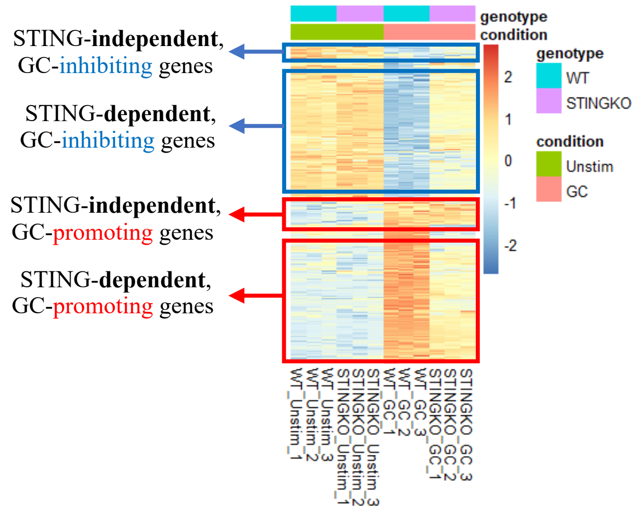


**c** **d**

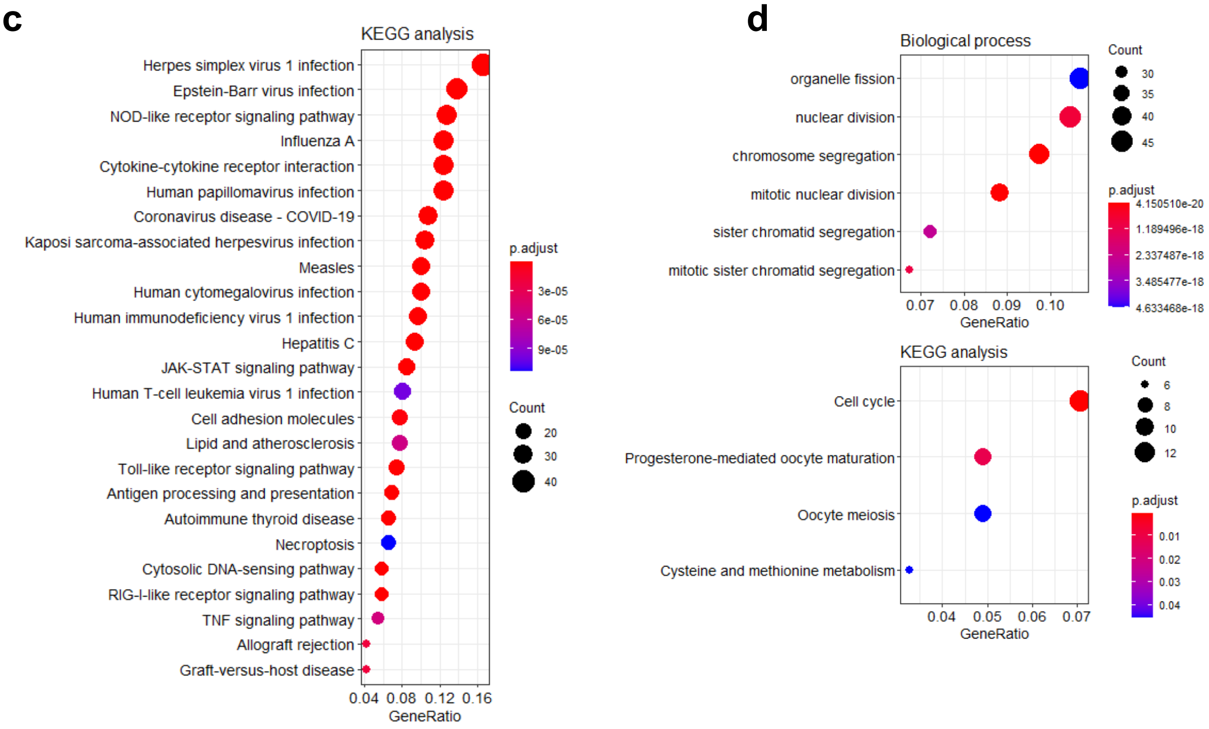


**Figure S5**

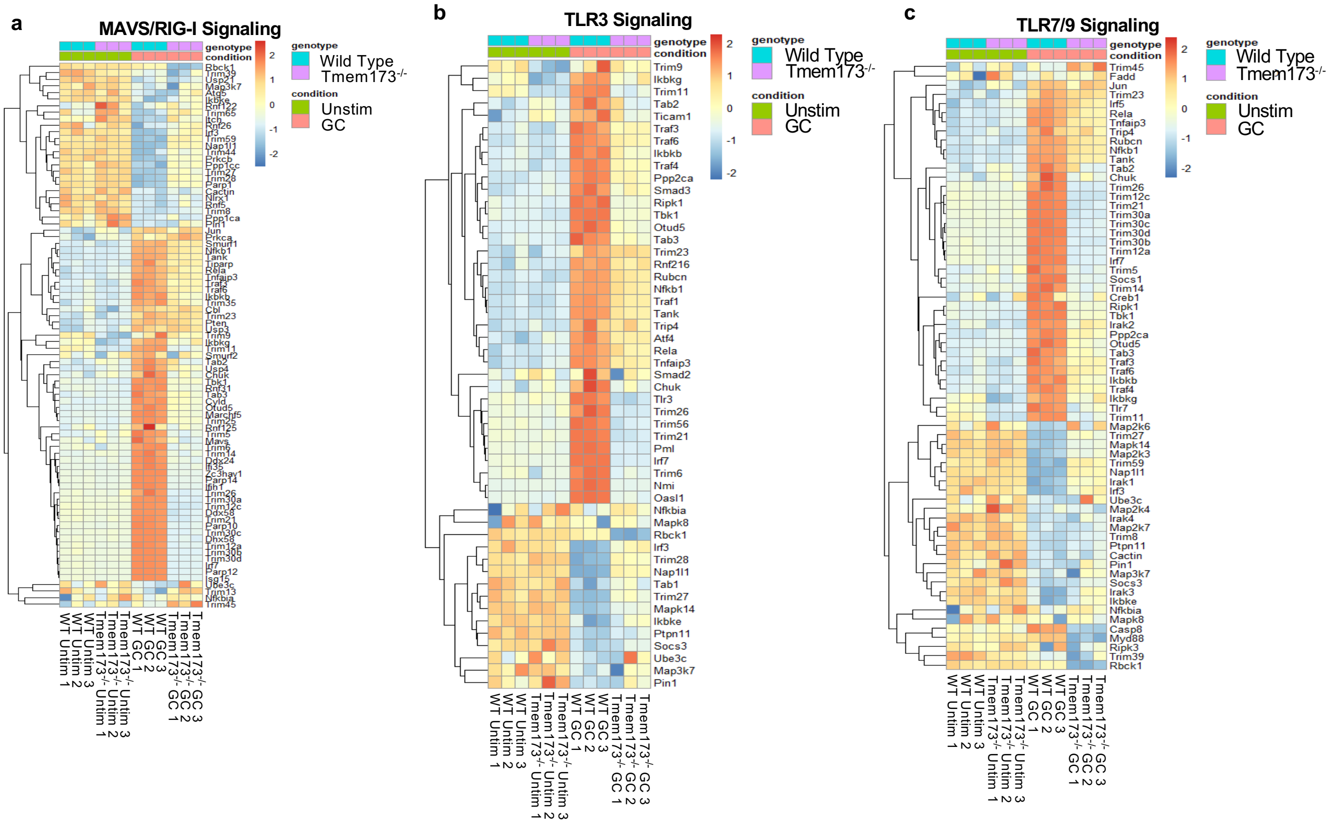
a
MAVS/RIG-I Signaling
b
TLR3 Signaling
c
TLR7/9 Signaling
genotype
condition
Wild Type
Tmem173-/-
Unstim
GC
2
1
0
-1
-2
Trim9
Ikbkg
Trim11
Tab2
Ticam1
Traf3
Traf6
Ikbkb
Traf4
Ppp2ca
Smad3
Ripk1
Tbk1
Otud5
Tab3
Trim23
Rnf216
Rubcn
Nfkb1
Traf1
Tank
Trip4
Atf4
Rela
Tnfaip3
Smad2
Chuk
Tlr3
Trim26
Trim56
Trim21
Pml
Irf7
Trim6
Nmi
Oasl1
Nfkbia
Mapk8
Rbck1
Irf3
Trim28
Nap1l1
Tab1
Trim27
Mapk14
Ikbke
Ptpn11
Socs3
Ube3c
Map3k7
Pin1
Trim45
Fadd
Jun
Trim23
Irf5
Rela
Tnfaip3
Trip4
Rubcn
Nfkb1
Tank
Tab2
Chuk
Trim26
Trim12c
Trim21
Trim30a
Trim30c
Trim30d
Trim30b
Trim12a
Irf7
Trim5
Socs1
Trim14
Creb1
Ripk1
Tbk1
Irak2
Ppp2ca
Otud5
Tab3
Traf3
Traf6
Ikbkb
Traf4
Ikbkg
Tlr7
Trim11
Map2k6
Trim27
Mapk14
Map2k3
Trim59
Nap1l1
Irak1
Irf3
Ube3c
Map2k4
Irak4
Map2k7
Trim8
Ptpn11
Cactin
Pin1
Map3k7
Socs3
Irak3
Ikbke
Nfkbia
Mapk8
Casp8
Myd88
Ripk3
Trim39
Rbck1
WT Unstim 1
WT Unstim 2
WT Unstim 3
Tmem173-/- Unstim 1
Tmem173-/- Unstim 2
Tmem173-/- Unstim 3
WT GC 1
WT GC 2
WT GC 3
Tmem173-/- GC 1
Tmem173-/- GC 2
Tmem173-/- GC 3

**Figure S6**

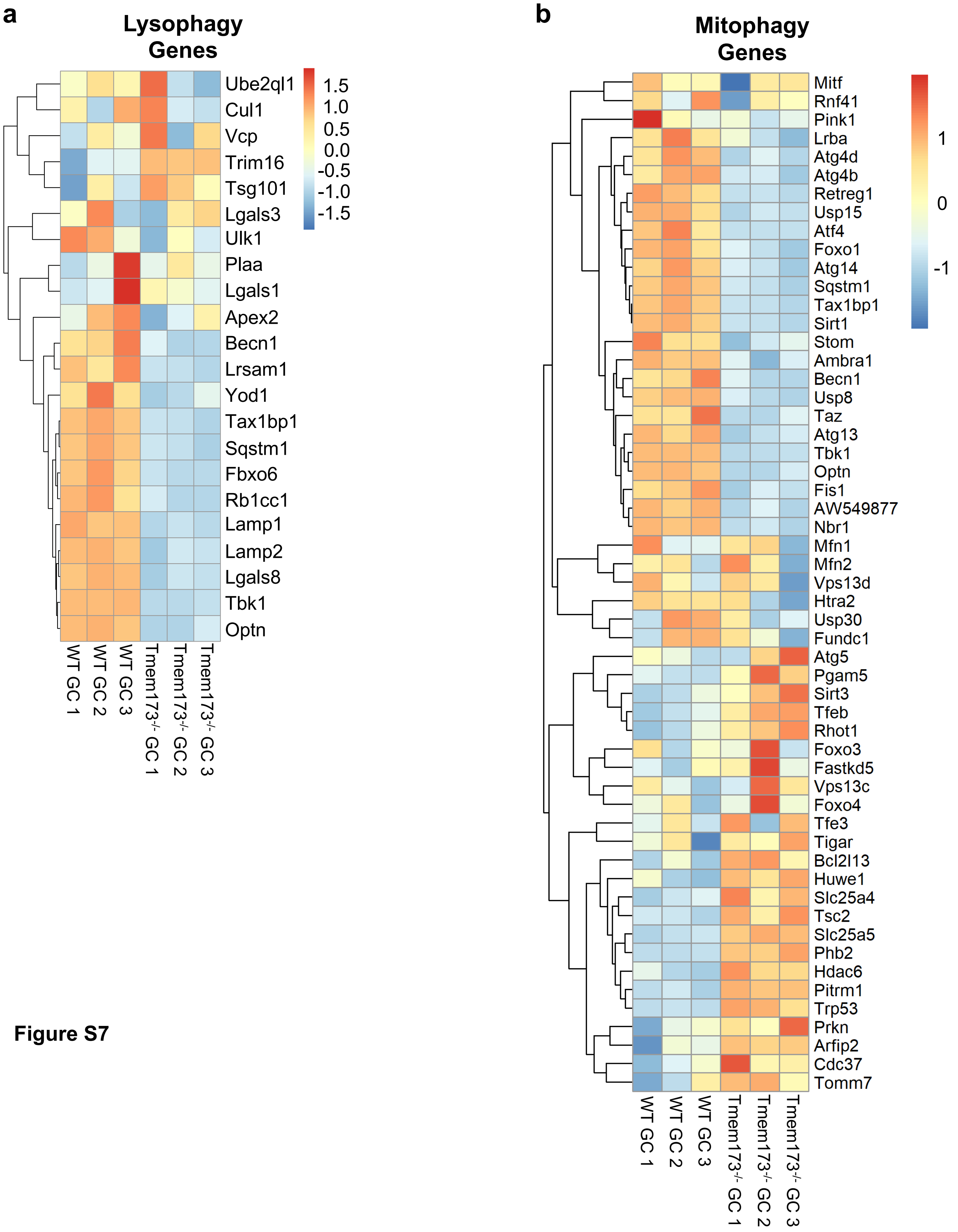
a
Lysophagy Genes
Ube2ql1
Cul1
Vcp
Trim16
Tsg101
Lgals3
Ulk1
Plaa
Lgals1
Apex2
Becn1
Lrsam1
Yod1
Tax1bp1
Sqstm1
Fbxo6
Rb1cc1
Lamp1
Lamp2
Lgals8
Tbk1
Optn
1.5
1.0
0.5
0.0
-0.5
-1.0
-1.5
WT GC 1
WT GC 2
WT GC 3
Tmem173-/- GC 1
Tmem173-/- GC 2
Tmem173-/- GC 3
b
Mitophagy Genes
Mitf
Rnf41
Pink1
Lrba
Atg4d
Atg4b
Retreg1
Usp15
Atf4
Foxo1
Atg14
Sqstm1
Tax1bp1
Sirt1
Stom
Ambra1
Becn1
Usp8
Taz
Atg13
Tbk1
Optn
Fis1
AW549877
Nbr1
Mfn1
Mfn2
Vps13d
Htra2
Usp30
Fundc1
Atg5
Pgam5
Sirt3
Tfeb
Rhot1
Foxo3
Fastkd5
Vps13c
Foxo4
Tfe3
Tigar
Bcl2l13
Huwe1
Slc25a4
Tsc2
Slc25a5
Phb2
Hdac6
Pitrm1
Trp53
Prkn
Arfip2
Cdc37
Tomm7
1
0
-1
WT GC 1
WT GC 2
WT GC 3
Tmem173-/- GC 1
Tmem173-/- GC 2
Tmem173-/- GC 3

**Figure S7**

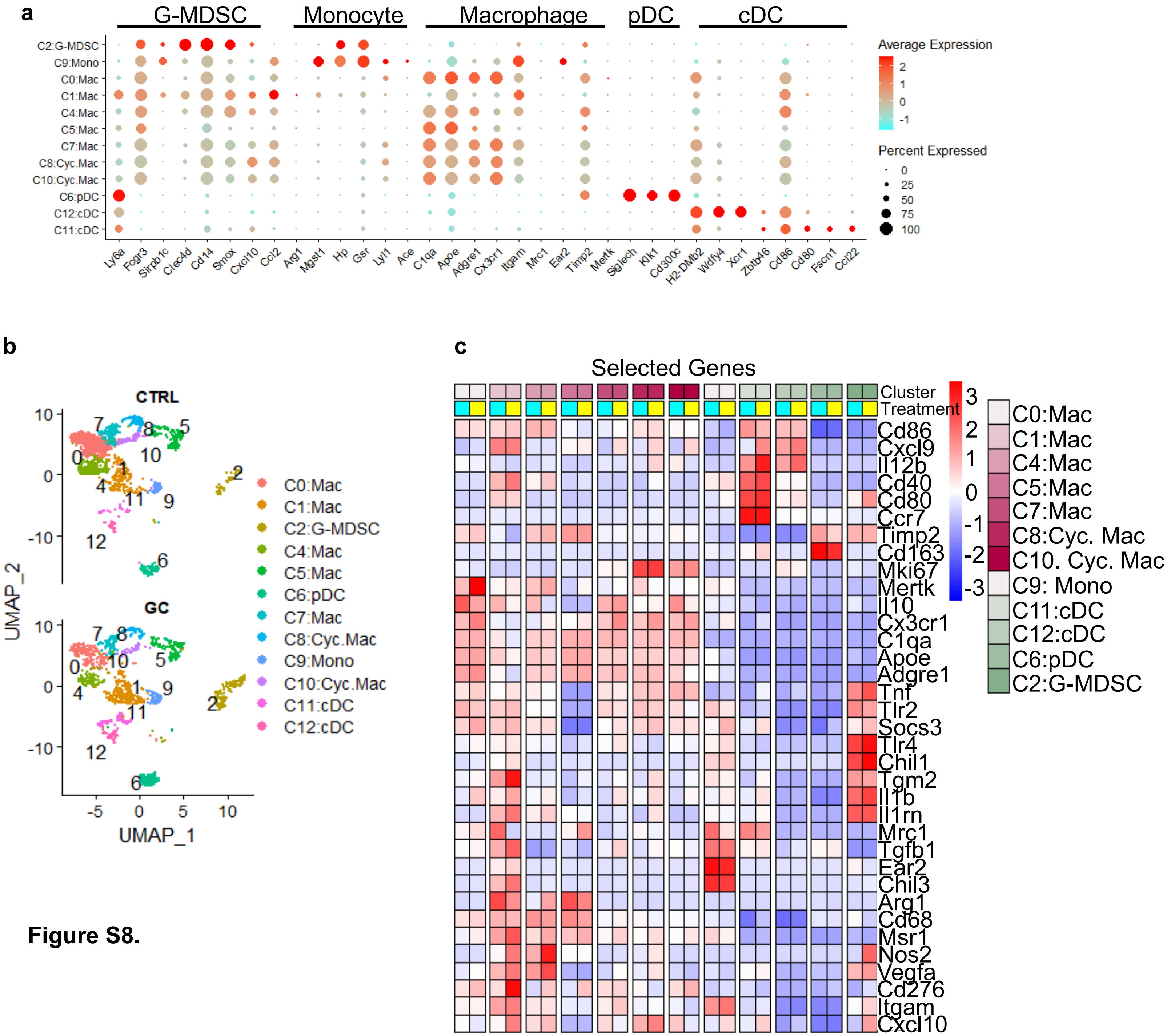
a
G-MDSC
Monocyte
Macrophage
pDC
cDC
C2:G-MDSC
C9:Mono
C0:Mac
C1:Mac
C4:Mac
C5:Mac
C7:Mac
C8:Cyc.Mac
C10:Cyc.Mac
C6:pDC
C12:cDC
C11:cDC
Ly6a
Fcgr3
Sirpb1c
Clec4d
Cd14
Smox
Cxcl10
Ccl2
Arg1
Mgst1
Hp
Gsr
Lyl1
Ace
C1qa
Apoe
Adgre1
Cx3cr1
Itgam
Mrc1
Ear2
Timp2
Mertk
Siglech
Klk1
Cd300c
H2-DMb2
Wdfy4
Xcr1
Zbtb46
Cd86
Cd80
Fscn1
Ccl22
Average Expression
2
1
0
-1
Percent Expressed
0
25
50
75
100
b
CTRL
GC
UMAP_2
UMAP_1
10
0
-10
-5
0
5
10
C0:Mac
C1:Mac
C2:G-MDSC
C4:Mac
C5:Mac
C6:pDC
C7:Mac
C8:Cyc.Mac
C9:Mono
C10:Cyc.Mac
C11:cDC
C12:cDC
c
Selected Genes
Cluster
Treatment
3
2
1
0
-1
-2
-3
Cd86
Cxcl9
Il12b
Cd40
Cd80
Ccr7
Timp2
Cd163
Mki67
Mertk
Il10
Cx3cr1
C1qa
Apoe
Adgre1
Tnf
Tlr2
Socs3
Tlr4
Chil1
Tgm2
Il1b
Il1rn
Mrc1
Tgfb1
Ear2
Chil3
Arg1
Cd68
Msr1
Nos2
Vegfa
Cd276
Itgam
Cxcl10
C0:Mac
C1:Mac
C4:Mac
C5:Mac
C7:Mac
C8:Cyc. Mac
C10. Cyc. Mac
C9: Mono
C11:cDC
C12:cDC
C6:pDC
C2:G-MDSC

**Figure S8.**
